# Changes in quantity and timing of foliar and reproductive phenology of tropical dry-forest trees under a warming and drying climate

**DOI:** 10.1101/2024.03.24.585819

**Authors:** Hao Ran Lai, Timothy Hill, Silvio Stivanello, Hazel M Chapman

## Abstract

1. Plant phenology drives population dynamics and forest productivity; it is also impacted by shifting environmental cues under climate change such as more prevalent drought. It is imperative to better understand how species and community phenology respond to climate change in leaf turnover and reproduction, both of which are required to integrate phenology into full life-cycle assessments.
2. However, relatively few studies to-date examined the quantity and timing of phenology simultaneously. We demonstrate that the simultaneous assessment of phenological quantity and timing across multiple organs reveals more nuanced and holistic insights into the consequences of climate change.
3. Extending a regression approach based on Fourier series, we decomposed the long-term (2004–2020) monthly leaf shedding, leaf flush, flowering and fruiting of 617 trees across 94 taxa at a Nigerian seasonally dry tropical forest into three periodic components—mean intensity, amplitude and phase—which respectively represents the total quantity, pulse concentration and peak timing of phenology. We then related each periodic component to warming minimum temperature and drying wet-season rainfall.
4. We found that climate explained more variation in phenological amplitudes (14–66%) compared to mean intensity and timing (6–49%). In drier years, more species (18%) shed leaves earlier (changing timing) or in more concentrated pulses (changing amplitude), while only a few (2%) shed leaves in greater total amounts (changing mean intensity). This combined with the decreased mean intensity of leaf flush imply a lower primary productivity as trees deployed fewer leaves for a shorter period. Some species (30%) produced fewer fruits despite no change or even increase in flowering; in a few species this could be explained by a shortened flowering period that limited pollination. At the community level, reproduction became more synchronous, potentially creating periods of scarcity for consumers.
5. **Synthesis:** Our findings highlight several contrasting yet complementary phenology– climate insights, indicating that assessments of forests’ climate resilience necessitate multiple aspects of phenology rather than a single performance indicator. The decline of leaf and fruit productions, as well as the temporal mismatches in leaf turnover and reproduction, will have cascading impacts on trophic interactions and nutrient cycling.

## Introduction

The phenology of tropical forest describes the temporal patterns of primary productivity and reproduction (Sakai, 2001), which have direct consequences on ecosystem processes such as nutrient cycling, multitrophic interaction, and species coexistence (Cleland & Wolkovich, 2024; Tang et al., 2016). However, the stability of these ecosystem processes are impacted by climate change as plants react by shifting the quantity and timing of reproduction and biomass turnover (Cleland et al., 2007; Iler et al., 2021). For example, more prevalent leaf shedding under the warmer and drier conditions in Amazonian forests has reduced trees’ ability to sequestrate carbon in living biomass (Janssen et al., 2021). Climate-induced changes in flowering and fruiting intensities across tropical forests have also impacted tree recruitment and consumer populations (Bush et al., 2020; Butt et al., 2015; Numata et al., 2022). As increasing drought and aridity weakens the role of tropical forests as carbon sinks (Corlett, 2016), it is imperative that we study phenological shifts as indicators of species’ and communities’ demographic responses to climate change (Hacket-Pain et al., 2024; Iler et al., 2021).

Across tropical forests, there is accumulating evidence of temperature and rainfall as key determinants of tree phenology (Sakai & Kitajima, 2019; Van Schaik et al., 1993). Water stress due to declining rainfall and rising temperature have been associated with greater quantity of leaf shedding (Janssen et al., 2021), as well as reduced quantity of leaf flush (Nomura et al., 2003), flowering (Butt et al., 2015; Lasky et al., 2016; Numata et al., 2022) and fruiting (Bush et al., 2020; C. A. Chapman et al., 2018); though the opposite or weaker relationships have also been found (Babweteera et al., 2018; C. A. Chapman et al., 2018; Pau et al., 2013; Wright & Calderón, 2006). In addition to shifting quantities, climate can also shift the *timing* of phenology (Chang-Yang et al., 2024; Chen et al., 2018). Changing rainfall and temperature patterns in recent decades have delayed flowering (Borchert, 1983), advanced leaf shedding (Reich & Borchert, 1984), and shortened leaf-flush intervals (Ho et al., 2024). However, evidence of climate effects on phenological timing in tropical forests remain sparse, because most studies, including the largest open-data initiative of global phenology (Hacket-Pain et al., 2022), analysed phenology as time-implicit total quantities (e.g., the weight of litterfall or abundance of individuals in flower summed or averaged over a period — usually a year) ignoring the timing shift of phenology *within* an annual cycle.

Extending on Newstrom et al. (1994), Sakai (2001) and Mendoza et al. (2018), we argue that it is important to examine both the quantity *and* timing of phenology, as they offer different insights into the ecosystem impacts of climate change. For example, complementing the quantity of annual leaf shedding with its timing not only informs us whether more nutrients are recycled into the soil, but also whether litterfall is decoupled temporally from other processes such as decomposition or seed germination. Here we focus on a regression approach that combines advances in Fourier analysis (Bush et al., 2017) and circular statistics (Davis et al., 2022) that decompose the periodic time series of phenology into three components: mean intensity, amplitude and phase (i.e., timing of peak) (Fidino & Magle, 2017; Nelson et al., 1979; Fig. 1).

**Figure 1:**
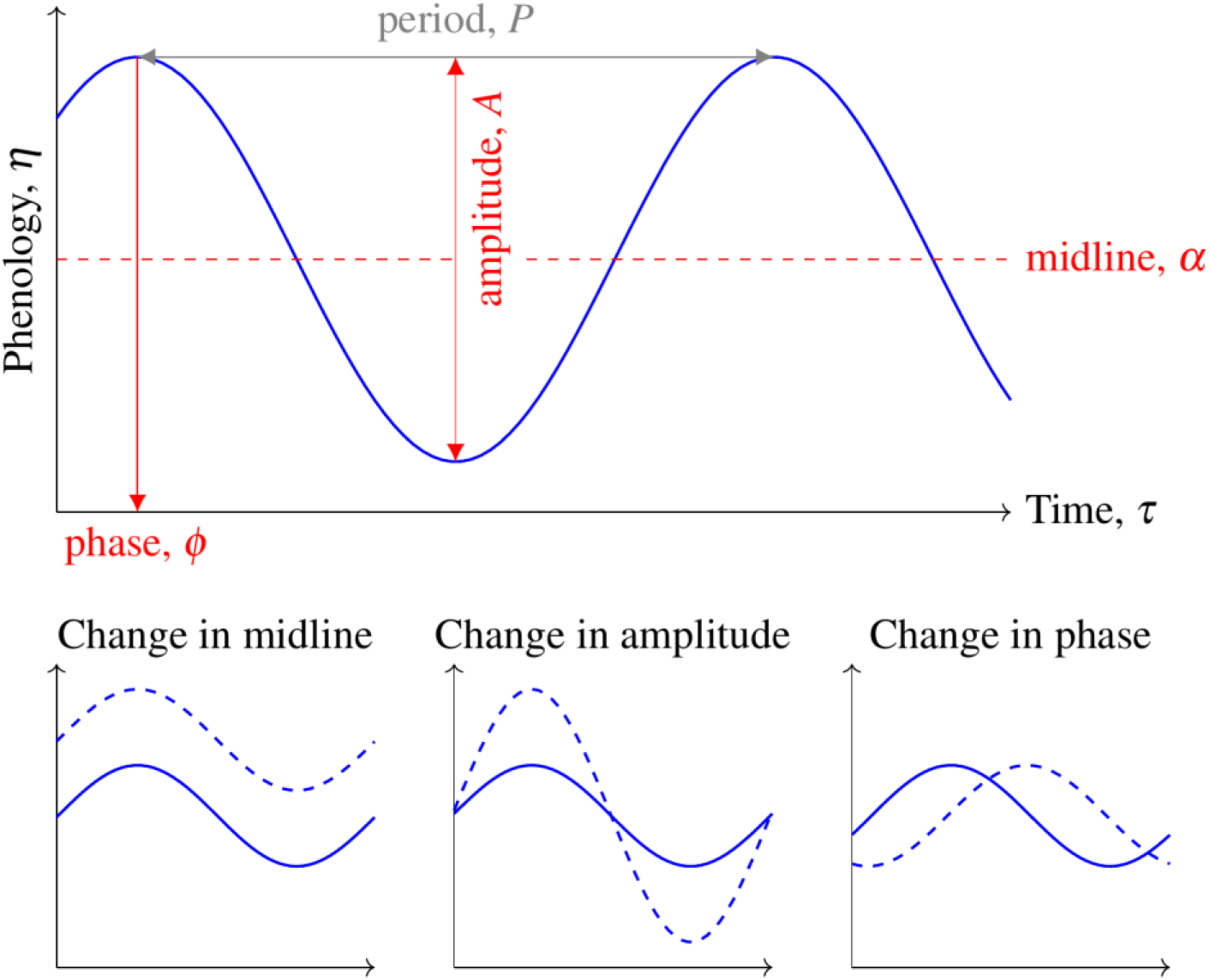
(Top panel) An example cosinor plot showing the basic components that correspond to Equation 1. Parameters in red are components that could be moderated by climate factors in this study. **(Bottom panels)** Examples illustrating how each cosinor components could be altered by climate factors.

The regression allows each periodic component to be hierarchically modelled as a function of climate to reveal different consequences on the ecosystem. Furthermore, we highlight the informative value in decomposing phenological quantity into mean intensity and *amplitude*; the latter can be interpreted as a more concentrated phenology within a shorter duration (this also implies that amplitude encompasses both quantity and timing, as in timespan). For example, an increase in amplitude leads to a more sudden pulse of leaf shedding *without changing its mean intensity*, therefore the *same total quantity* of leaves is shed within a narrower window of time (e.g., Ho et al. 2024), potentially leading to more transient ecosystem functioning (Sayer et al., 2024).

Additionally, a simultaneous examination of phenological quantity and timing, a pan-tropical synthesis of the effects of climate change on plant phenology also requires more complete data across organs. While there has been tremendous progress in tropical reproductive phenology (e.g., Hacket-Pain et al., 2022), foliar phenology could benefit from more long-term, species-specific data in the Tropics (Abernethy et al., 2018). This impedes the integration of phenological shifts into full life-cycle assessments, which are crucial to forecast population dynamics under climate change (Iler et al., 2021). This knowledge gap could be especially problematic for seasonally dry tropical forests (SDTFs), where leaf turnover is tightly regulated by strong rainfall and temperature seasonality (Murphy & Lugo, 1986; Van Schaik et al., 1993). SDTFs account for about 40% of the world’s tropical forests (Siyum, 2020) and may be more vulnerable to climate change due to their reliance on water availability (Aguirre-Gutiérrez et al., 2020). Nonetheless, a considerable phenological diversity exists among species in SDTFs (Lasky et al., 2016; Meir & Pennington, 2011; Singh & Kushwaha, 2016), providing a promising premise to learn the wide range of plant life-history strategies against past and future droughts.

From a pilot analysis of 47-year climate data in a Nigerian SDTF, we found that minimum temperature has risen by 0.8^∘^C while rainfall has become more erratic, with wet-season monthly rainfall decreased by about 10%; these changes were congruent to broader trends across tropical Africa (Bennett et al., 2021). To assess if climate change has altered the intensity, amplitude and timing of phenology, we modelled 17 years of monthly foliar and reproductive phenology—leaf shedding, leaf flush, flowering and fruiting—of 617 trees across 94 taxa as a function of rainfall and temperature. Additionally, to assess if SDTF species do have high phenological diversity with a wide range of drought adaptations as previously shown (K. Allen et al., 2017), we included the phenological responses to climate as species-specific slopes in the hierarchical regression. We show that the different responses of each periodic component provide a multifaceted interpretation on how climate influences phenology, and that a more consistent pattern may still emerge at the community level when phenological synchrony is calculated from apparently idiosyncratic changes in species-specific timings.

## Materials and Methods

### Study system

The study was carried out in the 46-km^2^ Ngel Nyaki Forest Reserve (7.06^∘^N, 11.1^∘^E) on the south west escarpment of the Mambilla Plateau in Taraba State, Nigeria (Fig. S1). Within the reserve, Ngel Nyaki forest is a 5.2-km^2^ stand of submontane forest on the steep slopes (1,600-m elevation) of an ancient volcano (J. D. Chapman & Chapman, 2001). The mean annual rainfall is approximately 1,800 mm, with most of the rain falling between April and October, followed by a six-month dry season. During the wet season, the forest can be covered in mist or fog for weeks on end, severely reducing irradiation (J. D. Chapman & Chapman, 2001). The mean annual temperature is 19^∘^C and the monthly mean maximum and minimum temperatures for the wet and dry seasons are 25.6 and 15.4^∘^C, and 28.1 and 15.5^∘^C, respectively.

Ngel Nyaki forest is relatively diverse for the Afromontane with at least 105 tree species from 47 families and 87 genera (Abiem et al., 2020). Forest species comprise a mix of Afromontane specialists (J. D. Chapman & Chapman, 2001), lowland forest and forest edge–grassland species. There is a gradient in species composition from forest core to edge, with edge species comprising more drought-tolerant grassland species (Abiem et al., 2020).

### Phenology data

Approximately 10 km^2^ of phenology transects were established in 2004. Under a systematic design, the transects are 500 m apart (Beck & Chapman, 2008), running east to west to criss-cross the forest and obtain a good representation of the community composition. Along the transects, 800 trees > 10 cm in diameter-at-breast-height (DBH), comprising 95 species were tagged, numbered and DBH measured. The number of trees per species ranged from 1 to 36 (median = 18.5). Tagged trees were chosen to ensure a representative sample of the forest composition including taxonomy, dispersal modes and flower types. During the monthly phenology monitoring from 2004 to 2020, trees are observed close-up, with binoculars when necessary. As an indicator of monthly leaf shedding, leaf flush, flowering and fruiting, the proportion of whole crown occupied by each phenological variable in a given tree is given an ordinal score between zero and four (0 = 0%, 1 = 1–25%, 2 = 26–50%, 3 = 51–75%, 4 = 76–100%) following Sun et al. (1996).

### Exploratory analyses of climate data

To provide context for climate-change impacts on phenology, we used a 47-year monthly time series from 1976–2022 from weather model-reanalysis data and in-situ observations. Rainfall was gathered from the Gembu State Government weather station (6^∘^ 41′ 13.08 N; 11^∘^ 17′ 33.48 E) 40 km from our study site. For ambient temperature, we used the 2-m air temperature product from the NASA ERA5-Land hourly data reanalysis dataset (Muñoz Sabater, 2019) with a 0.1^∘^ × 0.1^∘^horizontal resolution. Hourly values were converted to monthly means of daily minimum temperature.

Long-term climate trends in rainfall and minimum temperature were estimated the using linear regression, performed in `Python` v3.11.5 using the package `Seaborn` v0.12.2. Trends were fitted for individual months, as well as grouped months for dry (December, January and February), wet (June, July, August and September) and intermediate seasons (March, April, May, October and November). Additionally, we performed trend analysis using all months to examine overall changes. These exploratory analyses revealed significant increase and decline in all-month minimum temperature (hereafter “minimum temperature”) and wet-season rainfall, respectively. Therefore, we modelled monthly phenology as a function of minimum temperature and wet-season rainfall in the main regression as follows.

### Modelling framework for periodic phenology

To quantify the effects of climate change on the periodic components of tree phenology, we turn to a statistical approach known as cosine-vector or “cosinor” rhythmometry (Nelson et al., 1979). Cosinor is commonly used to model periodic time series that display regular cycles, and is generally defined nonlinearly as:

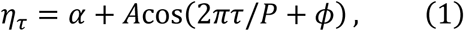

where a variable such as phenology, *η*_*τ*_, that varies over time *τ* is described by its midline *α* (mean intensity about which oscillation occurs), amplitude *A* (magnitude of fluctuation, or *concentration* of intensity as interpreted in this study) and phase *ϕ* (timing or starting point of the time series) for a given period *P* (e.g., a period of 12 months means that the time series has an annual cycle; see Fig. 1). It is possible to incorporate the potential effects of external factors, *X*, such as temperature on phenology by allowing them to influence the midline, amplitude and phase, for instance:

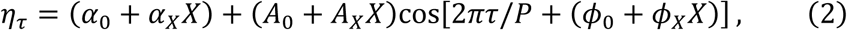

where *α*_*X*_, *A*_*X*_ and *ϕ*_*X*_ denote the effect of temperature on the midline, amplitude and phase, respectively. For example, positive values of *A*_*X*_ and *ϕ*_*X*_ would mean that warming temperature increases the amplitude and delays (right-shifts) the timing of phenology (Fig. 1).

In practice, however, fitting a statistical model based on Equation 2 could be challenging because of parameter non-identifiability, i.e., the parameters are often correlated causing model non-convergence. A solution lies in the rewriting of Equation 1 using the trigonometric identity cos(*u* + *v*) = cos(*u*)cos(*v*) − sin(*u*)sin(*v*), which is a common practice in cosinor modelling (Fidino & Magle, 2017):

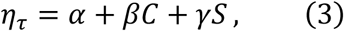

where *C* = cos(2*πτ*/*P*), *S* = sin(2*πτ*/*P*), *β* = *A*cos(*ϕ*) and *γ* = −*A*sin(*ϕ*). The reparameterised Equation 3 can then be fitted as a linear regression with intercept *α* and slopes *β* and *γ*, after which the original parameters can be recovered as 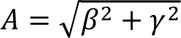 and *ϕ* = tan^−1^(−*γ*/*β*) (Nelson et al., 1979; Shumway & Stoffer, 2010). See Bingham et al. (1982) for additional sign correction for *ϕ*.

We could further include climate covariates into Equation 3:

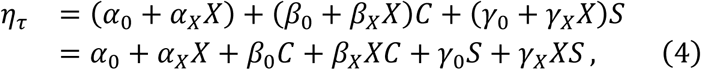

where *α*_0_ now serves as the intercept (midline); *α*_*X*_ as the main effect of climate (on midline); *β*_0_ and *γ*_0_ are the cosinor parameters under average climate condition (after centering the climate variable *X*); while *β*_*X*_and *γ*_*X*_are interactions between the cosinor and climate terms, i.e., the moderating effects of climate on amplitude and phase.

However, the computational efficiency of Equation 4 comes with a trade-off: we do not specify the relationship between climate, amplitude and phase directly. Instead, amplitude is now a function of climate as 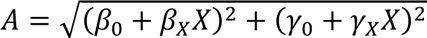, which when expanded contains the quadratic term *X*^2^and hence introduces unintended nonlinearity between amplitude and climate.

The same unintended nonlinearity also applies to the phase–climate relationship as *ϕ* = tan^−1^[−(*γ*_0_ + *γ*_*X*_*X*)/(*β*_0_ + *β*_*X*_*X*)]. In a pilot analysis, we fitted the direct parameterisation (Equation 2) but were unable to achieve model convergence. We also explored another option by fitting separate models: fitting Equation 1 first and then regressing the estimated coefficients as response variables to climate. However, we did not further pursue this approach because the model with phase as a response needed to account for its circular property (e.g., by using a von Mises distribution) and was also difficult to converge. Furthermore, fitting separate models do not propagate parameter uncertainties in the three periodic components. Therefore, we opted for the reparameterisation (Equation 4) and circumvented the unintended nonlinear climate effect using counterfactual comparisons (Arel-Bundock et al., 2024). Specifically, we calculated amplitude or phase under the minimum and maximum climate conditions (i.e., *X*_min_ and *X*_max_), then took the difference (i.e., *A* or *ϕ* under *X*_max_ minus that of *X*_min_) to represent the effect of climate of amplitude or phase.

### Model fitting

To apply the cosinor Equation 4 to our data, we fitted the ordinal scores *Y*_*pijnmt*_ of leaf shedding, leaf flush, flowering and fruiting of individual tree *i* of species *j* in transect *n*, month *m* and year *t* in a multivariate generalised linear mixed-effects model (GLMM) as cumulative processes with logit link:

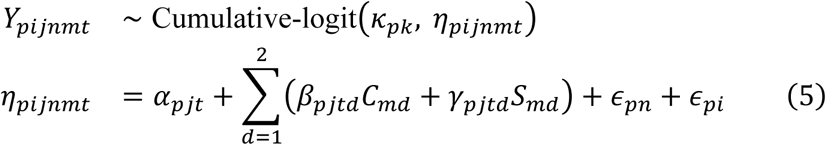

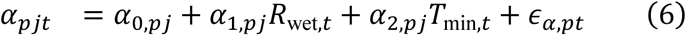

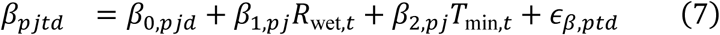

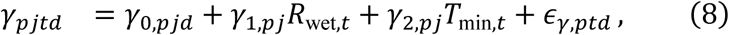

where subscript *p* denotes phenology of leaf shedding, leaf flush, flowering or fruiting. For each phenology, the model estimates an underlying latent, *continuous* variable *η* from which the five possible discrete ordinal scores *Y* ∈ {0,1, . . ., 4} were categorised and partitioned from four cutpoints, *κ*_*k*_, where *k* = 1,2, . . ., 4 (Bürkner & Vuorre, 2019, see Fig. S2 for an illustrated explanation). In other words, the model reflects that phenology is an underlying continuous quantity that was discretised by the field observers for logistical feasibility.

In the continuous linear predictor *η* (Equation 5), we began by including species-specific random intercepts, *α*_*pj*_, which model the midline for each species. Then we included the periodic terms *C*_*md*_ = cos(2*πm*/*P*_*d*_) and *S*_*md*_ = sin(2*πm*/*P*_*d*_), as well as the corresponding species-specific slopes *β*_*pjtd*_ and *γ*_*pjtd*_ to model species’ phenological amplitudes and phases. Note that we included two sets of periodic terms (hence the summation up to *d* = 2) to estimate the relative signals in annual (*P*_*d*=1_ = 12) and sub-annual (*P*_*d*=2_ = 6) cycles for each species. This is essentially a Fourier transformation to break down monthly phenologies into two dominant cycles (Fidino & Magle, 2017), and we chose the 12- and 6-month cycles based on Bush et al. (2017) from the same biome.

Next, we inferred the effects of annual climate factors—wet-season rainfall *R*_wet_ and minimum temperature *T*_min_—on phenology. In a pilot analysis, we included both current- and previous-year climate to examine lag effects and if species with phenologies that peak in earlier months would respond more to previous-than to current-year climate. We found little evidence that previous-year climate was as important as current-year climate (Fig. S3) and therefore opted to only use current-year climate factors for simplicity. In the midline submodel (Equation 6), *α*_0,*pj*_ denotes the species-specific midline under the average climate condition (after centering *R*_wet,*t*_ and *T*_min,*t*_), while *α*_1,*pj*_ and *α*_2,*pj*_ are the effects of wet-season rainfall and minimum temperature on species’ midlines, respectively. We further included the year-specific random error *ε*_*α*,*pt*_to account for residual variation in species’ midlines unexplained by climate. We also specified similar submodels for the other two cosinor parameters, *β*_*pjtd*_ and *γ*_*pjtd*_, to infer the effects of climate on species’ annual and sub-annual amplitudes and phases (Equations 7 and 8). It is important to note that the annual climate covariates (as well as the year-specific random errors *ε*_*α*,*pt*_, *∈*_*β*,*pt*_, and *∈*_*γ*,*pt*_) address our study goal by accounting for potential non-stationary in the time series, i.e., by allowing each year to have different phenological midline, amplitude and phase (note the *t* subscript in the year-varying *α*, *β* and *γ* terms). Lastly, we included transect- and individual-random intercepts, *∈*_*pn*_ and *∈*_*pi*_, to account for spatial and among-tree non-independence (Equation 5).

Prior to modelling, we selected living tree individuals with at least 10 years of records and which had no observation gap for > 3 months, were not too tall for reliable measurement, and did not have constant phenology during the study period. We also grouped all *Ficus* spp. into a single taxonomic unit. This resulted in a total of 117,964 observations from 617 trees across 94 taxa, 17 transects and 17 years. The climate variables were centred and scaled to unit standard deviation to promote model convergence.

The model was fitted with the `brms` package v2.18.3 (Bürkner, 2017) in `R` v4.2.1 (R Core Team, 2022). Four chains of Hamiltonian Monte Carlo (HMC) iterations were run, each with 2,000 iterations and the first 1,000 samples as warmup. We used the default weakly informative priors for all parameters and a target average acceptance probability of 0.95 in `brms`. Chain convergence was assessed visually using trace plots and the Gelman–Rubin diagnostic *R*^ < 1.05.

### Community synchrony

To examine how climate-induced shifts in species’ phenological timing lead to changes in community synchrony, we calculated the circular standard deviation (SD) of the estimated values of phases *ϕ* across climate gradients of *R*_wet_ and *T*_min_ following Bush et al. (2017) using the `R` package `circular` (Agostinelli & Lund, 2024). The SD was inverted to precision so that higher values indicate greater synchrony. We acknowledge that there are alternative metrics to calculate synchrony from the whole, rather than only the timing, of time series (e.g., Gouhier & Guichard, 2014) but opted to follow Bush et al. (2017) since our modelling framework is more compatible with theirs.

## Results

### Overall temporal trends in climate and phenology

From 1976 to 2022, there was no clear annual rainfall trend in the Mambilla plateau where our forest is situated (Fig. S4). When broken down into seasons, however, an average month in the dry and wet seasons had become wetter by 18 mm and drier by 30 mm over the past 46 years, respectively (Figs 2a and S4). During the same period, minimum temperature had increased significantly by 0.017^∘^C per year, or 0.8^∘^C in total (Fig. 2a and S5).

**Figure 2:**
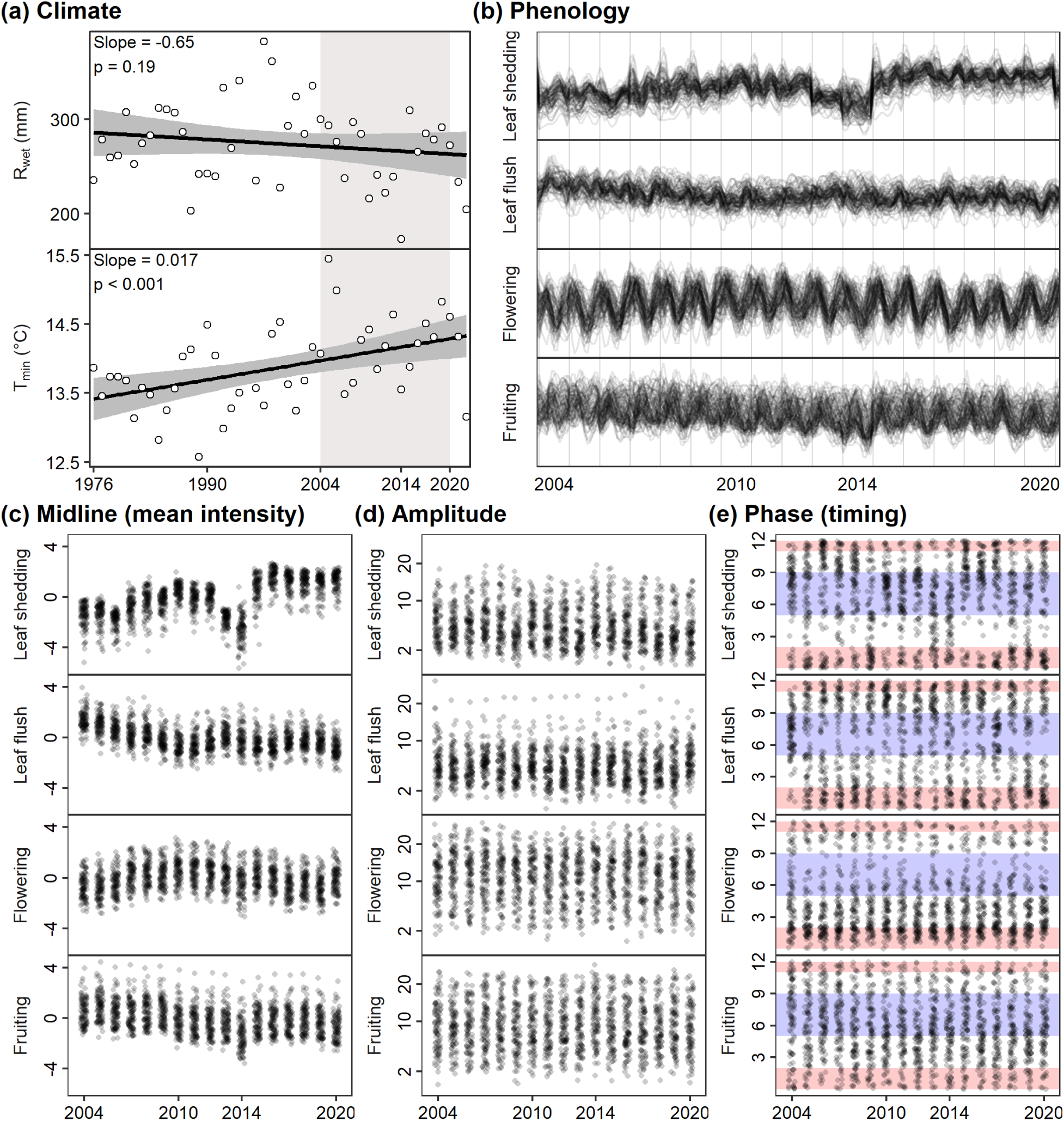
Temporal trends in climate and phenology. **(a)** Annual changes in monthly wet-season rainfall (*R*_wet_) and minimum temperature (*T*_min_) in the Mambilla plateau, where our study site was situated, over nearly half a century. Thick black lines and grey shaded regions are fitted trends and 90% confidence intervals, respectively. Grey box highlights the period in which we collected phenological data. **(b)** Model-fitted values of species-specific monthly phenology from 2004 to 2020 (thin black lines). **(c**–**d)** Temporal trends in the midline (mean intensity), amplitude and phase (timing) of phenology. Points represent species-specific posterior median estimates and are jittered along the X-axis for visualisation. Note the square-root scale on the Y axis in (d). Blue and red shades in (e) denote wet and dry seasons, respectively.

Within this warming and drying period, we recorded monthly foliar and reproductive phenologies from 2004 to 2020 and modelled the changes in their midline (mean intensity), amplitude and timing (Fig. 2b). Almost all species exhibited annual cycles in reproduction, while a few more species exhibited sub-annual cycles in foliar phenology (Fig. S6). Over time, there were considerable fluctuations and but overall an increase in the mean intensity of leaf shedding, whereas leaf flush and fruiting had overall decreased slightly in mean intensities (Fig. 2c). The trends in the amplitudes and timings of phenology were less consistent; there was a slight decrease in leaf-shedding amplitudes (Fig. 2d) and considerable fluctuations in the timing of foliar phenology across the years (Fig. 2e). Overall, the timing of fruiting was regularly spaced throughout a year, but there tended to be a gap in flowering towards the end of wet season, a gap in leaf shedding during the intermediate seasons, as well as a slight gap in leaf flush in the middle of wet season (Figs 2e and S7).

### Climate effects on aspects of phenology

To explain the temporal changes in phenology, we modelled phenology as a function of wet-season rainfall (*R*_wet_) and minimum temperature (*T*_min_) and found that, across periodic components, climate explained the greatest amount of temporal variation in amplitudes (14– 66%), followed by midlines (18–49%) and then phases (6–20%; Fig. S8). Across organs, climate explained the most temporal variation in fruiting and the least in leaf flush. For example, the year 2014 experienced the driest wet season during our study, which corresponded to less intense but earlier leaf shedding, less intense fruiting, and varied amplitudes in all phenologies (Figs 2c–e).

Specifically, the effect of climate on midlines had a more consistent direction than the effects on amplitudes and phases. The midline of leaf shedding, leaf flush and fruiting of most species decreased with drying *R*_wet_, while the midline of flowering decreased with warming *T*_min_ (left column of Fig. 3 and Figs S9–S10). The phenological amplitudes of many species were more strongly influenced by climate, but the huge interspecific variation and mixed signs in climate effects resulted in an overall more neutral change at the community level (middle column of Fig. 3 and Figs S11–S12). There were also some species-specific changes in the phases or timing of phenology with climate, but they tended to have much greater uncertainties (right column of Fig. 3 and Figs S13–S14).

**Figure 3:**
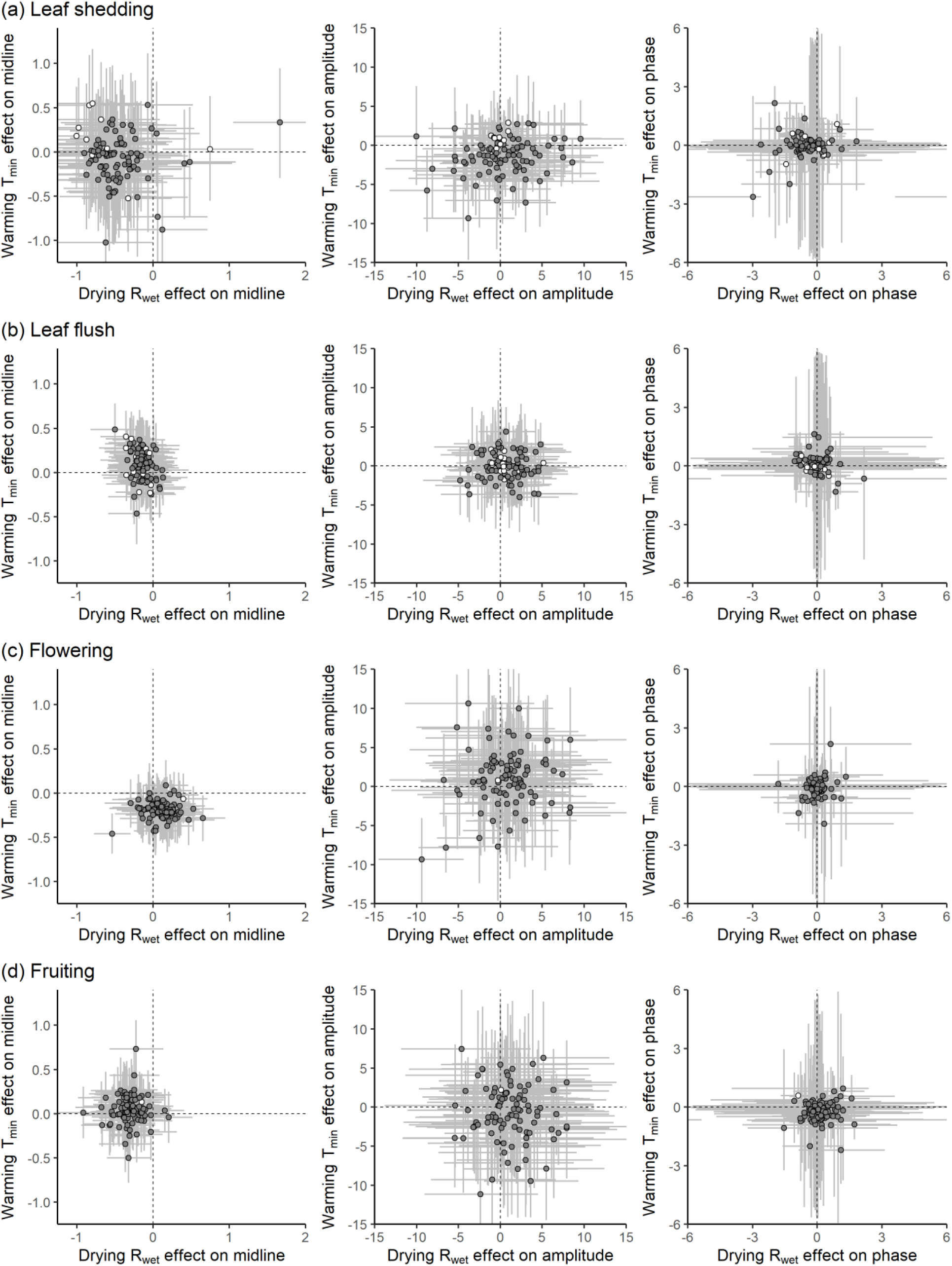
The effects of decreasing annual wet-season rainfall (X-axes, *R*_wet_) and increasing minimum temperature (Y-axes, *T*_min_) on the three aspects of phenology— midline (1st column), amplitude (2nd column), and phase (3rd column)—for leaf shedding (a), leaf flush (b), flowering (c), and fruiting (d). Symbols denote the posterior median of species, while error bars are the 89% credible intervals of posterior distribution. Grey and white circles represent species with predominantly annual and subannual phenology, respectively.

### Climate effects on community synchrony

The community synchronies in reproductive phenology were more strongly associated with climate than foliar phenology (Fig. 4). With decreasing *R*_wet_ and increasing *T*_min_, both flowering and fruiting became more synchronous, while the synchrony of foliar phenology did not change strongly except for a slight decrease in leaf-shedding synchrony.

**Figure 4:**
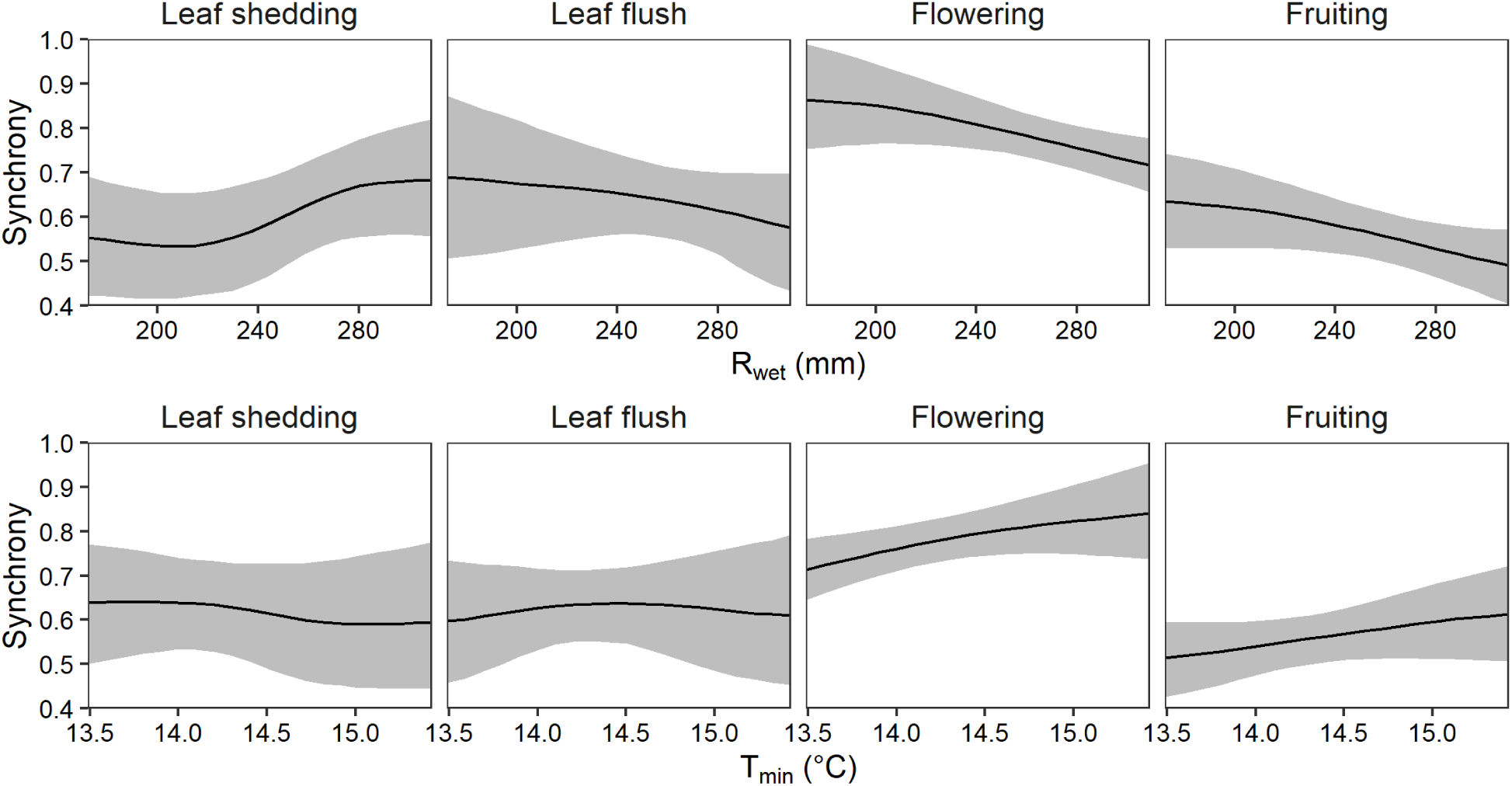
Community synchrony of phenology (calculated as the standard deviation of phases across species following Bush et al., 2017) along gradients of wet-season rainfall, *R*_wet_ **(top panels)** and minimum temperature, *T*_min_ **(bottom panels)**. Line and shaded regions denote the posterior median and the 89% credible intervals.

## Discussion

Over 17 years in a seasonally dry tropical forest (SDTF) in Nigeria, we observed the foliar and reproductive phenology of 94 tree taxa, which mostly exhibited annual cycles in leaf shedding, flowering, and fruiting. This corroborates with previous studies in African forests (Adamescu et al., 2018; Bush et al., 2017; Ouédraogo et al., 2020), where periodic phenology evolved under strong environmental cues across distinct wet–dry seasons (Borchert 1983; van Schaik, 1993; Sakai, 2001; Daïnou 2012). Leaf flush, however, displayed sub-annual cycles in more species and indicates an adaptive strategy to capitalise intermittent rainfall events that characterise our dry season (Lieberman and Lieberman, 1984). As observed in other African forests (Chapman et al 1999; Adamescu, 2018), our species varied considerably in the quantity and timing of foliar and reproductive phenologies, reflecting their survival strategies to seasonal abiotic and biotic cues (Daïnou et al 2012; Singh and Kushwaha 2005; Williams et al 2008; Lensky 2016) and roles in providing food supply for frugivores and pollinators spread across the year (Chapman et al 2018). More importantly, we detected huge interspecific and among-organ variations in the phenological timing and quantity (which we decomposed into mean intensity and amplitude) in response to decreasing wet-season rainfall (*R*_wet_) and increasing minimum temperature (*T*_min_).

Consequently, the driest year of our study (2014) displayed less intense and earlier leaf shedding, less intense fruiting and varied amplitudes in all phenologies across species.

### Phenological amplitude reveal more nuanced insights than intensity or timing alone

One of the emergent patterns from the phenology–climate relationships was changing deciduousness. Whereas previous tropical studies have demonstrated increased deciduousness with leaf-shedding intensity or timing under drier conditions (Borchert et al., 2002; Condit et al., 2002; Edwards et al., 2018; Janssen et al., 2021; Wu et al., 2021), we show that increased deciduousness could also result from increased amplitude. In our study, amplitude was more sensitive to climate change than timing and intensity. Twenty percent of species in total showed strong increases in deciduousness with drying *R*_wet_ but in different periodic components: 2% shedded more leaves due to increased mean intensity, 8% shedded leaves earlier due to advanced timing, 6% had more concentrated leaf shedding due to increased amplitude, while 4% had more concentrated and earlier leaf shedding due to changes in both timing and amplitude. Therefore, most of our species with increased deciduousness did not shed more leaves in total under drier conditions, instead they tended to shed the same amount of leaves in a more concentrated pulse within a shorter window of time. This phenomenon could be due to reduced leaf flush in a previously dry year (this study and Ho et al., 2024), so that there was less foliar biomass to begin with, highlighting the importance of considering the lag and cumulative effects of consecutive droughts (Wunderling et al., 2022). The combination of reduced leaf flush followed by more abrupt and earlier leaf shedding implies that trees hold onto fewer leaves for a shorter period; there could be cascading forest die-off if droughts continue to exacerbate such a feedback (C. D. Allen et al., 2015).

Including amplitude in the analysis also helped reconcile an apparent conflict between the results of flowering and fruiting. For example, *Anthonotha noldeae* is a widespread tree species in our forest with increased mean flowering intensity, but decreased mean fruiting intensity, in drier years. A possible explanation for the mismatch between flower and fruit quantities is the increased flowering amplitude of *A. noldeae* in drier years; this indicates that the species flowered for a shorter duration, thus providing less opportunity for bird pollinators that are already rare to visit the flowers even when they pulsed in higher numbers (Beavon & Chapman, 2011). When the phenology of a widespread species that produces hyperabundant floral and fruit resources is affected by climate change, there could be cascading effects across the food web (Renner & Zohner, 2018). The fruiting of another common species, *Carapa oreophila*, not only had reduced mean intensity but also increased amplitude in drier years; this meant fewer fruits in total and for a shorter period. Such a compounding decline in fruit availability may drastically shift the behavior of being a key seed disperser—the African giant pouched rat (*Cricetomys* sp.)—from a scatterhoarding mutualist to a seed predator (Yadok et al., 2019), which would then feedback to a population decline of the primary producer.

### Phenological mean intensity had more consistent climate responses

Compared to amplitude, the mean intensity of phenology had a more unidirectional response to climate, especially *R*_wet_. In drier years, mean intensities tended to decrease for most species in all phenologies except flowering. As most species produce less litter, fewer leaves and fewer fruits in total, this indicates a lower primary productivity overall in drier conditions as previous studies suggested (Castro et al., 2018; Laan-Luijkx et al., 2015; Xu et al., 2019). Although the reduced leaf shedding during drought may suggest that trees were holding onto their leaves as a form of drought tolerance, there might be alternative mechanisms at play, such as the dust-laden harmattan winds of the dry season (Jenik & Hall, 1966) or aerial particulates from fires that could reduce irradiation to a point where evapotranspiration is less exacerbated by drought (Wright & Cornejo, 1990). The decreased mean fruiting intensities in drier years despite the opposite trend in flowering suggests poorer floral quality (Kuppler & Kotowska, 2021), resulting in higher rates of fruit abortion (Bawa & Webb, 1984). Surplus flower production may also be a strategy to satiate predispersal fruit and seed predators in harsher environments (Stephenson, 1981), but support for this hypothesis remains weak (Jeffs et al., 2018). In drier years, 30 of our species had strong decline in mean fruiting intensity with no change or even an increase in mean flowering intensity; this could be due to pollination mismatch by shortened flowering duration (increased amplitudes) in three species (*A. noldeae*, *Diospyros monbuttensis* and *Macaranga occidentalis*) or delayed flowering in two species (*Isolona deightonii* and *Sterculia trigantha*).

For the remaining species, more research is urgently needed to explain such a decoupling between mean flower and fruit quantities under drought conditions, especially when five species (including *I. deightonii*) are important fruit providers for the endangered Nigeria–Cameroon chimpanzee (*Pan troglodytes ellioti*).

### Phenological phase had uncertain climate responses but led to more consistent synchrony trends

There were high uncertainties around the effects of climate on the timing of species-level phenology but they lined up to more consistent trends in community synchrony, especially for reproduction. Flowering and fruiting were more synchronous in drier years, indicating a tighter schedule around the onset of wet season for trees to reproduce. A more synchronous flower and fruit production could also create periods of scarcity for pollinators and frugivores (Renner & Zohner, 2018), thereby increasing competition among consumers due to behavioural changes or fewer opportunities to temporally partition resources (Ovaskainen et al., 2013). In contrast, foliar synchrony changed more mildly across climate gradients, though leaf shedding were slightly less synchronous in drier years possibly due to strong bidirectional changes in the timing of leaf shedding among species. Such a more evenly distributed leaf-litter deposition throughout the year could alter the rate of nutrient cycling, primary productivity and window for seed germination (Sayer, 2006; Wood et al., 2009).

### Conclusions

Using a long-term community dataset of multi-organ phenology, we show that examining both the quantity and timing of phenology contributed to a more nuanced and complete understanding about the consequences of climate change on ecosystem functioning. There was a high diversity of phenology–climate relationships across the 94 taxa, yet a few general patterns emerged: interspecific phenological responses to climate change were more consistent in terms of total quantity and less consistent in terms of timing (but more consistent when viewed as synchrony at the community level). Furthermore, the duration or concentration of phenology potentially reconciles some of the disparate trends in the mean quantity of phenology between organs. In our study, maintaining or increasing flower quantity did not guarantee that fruit quantity would not decline, which could be explained by shortened flowering durations. If these ecological events continue to take place within shorter windows of time, then plant-supported ecosystem functions will become more ephemeral and mismatched against consumer phenology, even when total productions remain unchanged. Lastly, the modelling framework applied here could be vastly improved by the inclusion of life-history traits (Lasky et al., 2016) and phylogeny (Davis et al., 2022) towards a more generalised and mechanistic understanding.

## References

Abernethy, K., Bush, E. R., Forget, P. M., Mendoza, I., & Morellato, L. P. C. (2018). Current issues in tropical phenology: a synthesis. Biotropica, 50(3), 477–482. 10.1111/btp.12558

Abiem, I., Arellano, G., Kenfack, D., & Chapman, H. (2020). Afromontane Forest Diversity and the Role of Species Distribution. Diversity, 12(30), 1–19. www.mdpi.com/journal/diversity

Adamescu, G. S., Plumptre, A. J., Abernethy, K. A., Polansky, L., Bush, E. R., Chapman, C. A., Shoo, L. P., Fayolle, A., Janmaat, K. R. L., Robbins, M. M., Ndangalasi, H. J., Cordeiro, N. J., Gilby, I. C., Wittig, R. M., Breuer, T., Hockemba, M. B. N., Sanz, C. M., Morgan, D. B., Pusey, A. E., … Beale, C. M. (2018). Annual cycles are the most common reproductive strategy in African tropical tree communities. Biotropica, 50(3), 418–430. 10.1111/btp.12561

Agostinelli, C., & Lund, U. (2024). R package `circular`: Circular statistics (version 0.5-1). https://CRAN.R-project.org/package=circular

Aguirre-Gutiérrez, J., Malhi, Y., Lewis, S. L., Fauset, S., Adu-Bredu, S., Affum-Baffoe, K., Baker, T. R., Gvozdevaite, A., Hubau, W., Moore, S., Peprah, T., Ziemińska, K., Phillips, O. L., & Oliveras, I. (2020). Long-term droughts may drive drier tropical forests towards increased functional, taxonomic and phylogenetic homogeneity. Nature Communications, 11(1), 1–10. 10.1038/s41467-020-16973-4

Allen, C. D., Breshears, D. D., & McDowell, N. G. (2015). On underestimation of global vulnerability to tree mortality and forest die-off from hotter drought in the Anthropocene. Ecosphere, 6(8), 1–55. 10.1890/ES15-00203.1

Allen, K., Dupuy, J. M., Gei, M. G., Hulshof, C., Medvigy, D., Pizano, C., Salgado-Negret, B., Smith, C. M., Trierweiler, A., Van Bloem, S. J., Waring, B. G., Xu, X., & Powers, J. S. (2017). Will seasonally dry tropical forests be sensitive or resistant to future changes in rainfall regimes? Environmental Research Letters, 12(2). 10.1088/1748-9326/aa5968

Arel-Bundock, V., Greifer, N., & Heiss, A. (2024). How to interpret statistical models using marginaleffects in R and Python. Journal of Statistical Software, 111(9), 1–32. 10.18637/jss.v111.i09.

Babweteera, F., Plumptre, A. J., Adamescu, G. S., Shoo, L. P., Beale, C. M., Reynolds, V., Nyeko, P., & Muhanguzi, G. (2018). The ecology of tree reproduction in an African medium altitude rain forest. Biotropica, 50(3), 405–417. 10.1111/btp.12563

Bawa, K. S., & Webb, C. J. (1984). Flower, Fruit and Seed Abortion in Tropical Forest Trees: Implications for the Evolution of Paternal and Maternal Reproductive Patterns. American Journal of Botany, 71(5), 736. 10.2307/2443371

Beavon, M. A., & Chapman, H. M. (2011). Andromonoecy and high fruit abortion in Anthonotha noldeae in a West African montane forest. Plant Systematics and Evolution, 296(3-4), 217–224. 10.1007/s00606-011-0488-1

Beck, J., & Chapman, H. (2008). A population estimate of the Endangered chimpanzee Pan troglodytes vellerosus in a Nigerian montane forest: Implications for conservation. Oryx, 42(3), 448–451. 10.1017/S0030605308001397

Bennett, A. C., Dargie, G. C., Cuni-Sanchez, A., Mukendi, J. T., Hubau, W., Mukinzi, J. M., Phillips, O. L., Malhi, Y., Sullivan, M. J. P., Cooper, D. L. M., Adu-Bredu, S., Affum-Baffoe, K., Amani, C. A., Banin, L. F., Beeckman, H., Begne, S. K., Bocko, Y. E., Boeckx, P., Bogaert, J., … Lewis, S. L. (2021). Resistance of African tropical forests to an extreme climate anomaly. Proceedings of the National Academy of Sciences of the United States of America, 118(21), 1–12. 10.1073/pnas.2003169118

Bingham, C., Arbogast, B., Cornelissen Guillaume, G., Lee, J. K., & Halberg, F. (1982). Inferential statistical methods for estimating and comparing cosinor parameters. Chronobiologia, 9(4), 397–439.

Borchert, R. (1983). Phenology and Control of Flowering in Tropical Trees. Biotropica, 15(2), 81. 10.2307/2387949

Borchert, R., Rivera, G., & Hagnauer, W. (2002). Modification of vegetative phenology in a tropical semi-deciduous forest by abnormal drought and rain. Biotropica, 34(1), 27–39. 10.1111/j.1744-7429.2002.tb00239.x

Bürkner, P. C. (2017). brms: An R package for Bayesian multilevel models using Stan. Journal of Statistical Software, 80(1), 1–28. 10.18637/jss.v080.i01

Bürkner, P. C., & Vuorre, M. (2019). Ordinal Regression Models in Psychology: A Tutorial. Advances in Methods and Practices in Psychological Science, 2(1), 77–101. 10.1177/2515245918823199

Bush, E. R., Abernethy, K. A., Jeffery, K., Tutin, C., White, L., Dimoto, E., Dikangadissi, J. T., Jump, A. S., & Bunnefeld, N. (2017). Fourier analysis to detect phenological cycles using long-term tropical field data and simulations. Methods in Ecology and Evolution, 8(5), 530–540. 10.1111/2041-210X.12704

Bush, E. R., Jeffery, K., Bunnefeld, N., Tutin, C., Musgrave, R., Moussavou, G., Mihindou, V., Malhi, Y., Lehmann, D., Ndong, J. E., Makaga, L., & Abernethy, K. (2020). Rare ground data confirm significant warming and drying in western equatorial Africa. PeerJ, 2020(4), 1–29. 10.7717/peerj.8732

Butt, N., Seabrook, L., Maron, M., Law, B. S., Dawson, T. P., Syktus, J., & Mcalpine, C. A. (2015). Cascading effects of climate extremes on vertebrate fauna through changes to low-latitude tree flowering and fruiting phenology. Global Change Biology, 21(9), 3267–3277. 10.1111/gcb.12869

Castro, S. M., Sanchez-Azofeifa, G. A., & Sato, H. (2018). Effect of drought on productivity in a Costa Rican tropical dry forest. Environmental Research Letters, 13(4). 10.1088/1748-9326/aaacbc

Chang-Yang, C. H., Chiang, P. H., Wright, S. J., Hsieh, C. F., & Sun, I. F. (2024). Proximate cues of flowering in a subtropical rain forest. Biotropica, 56(1), 78–89. 10.1111/btp.13282

Chapman, C. A., Valenta, K., Bonnell, T. R., Brown, K. A., & Chapman, L. J. (2018). Solar radiation and ENSO predict fruiting phenology patterns in a 15-year record from Kibale National Park, Uganda. Biotropica, 50(3), 384–395. 10.1111/btp.12559

Chapman, J. D., & Chapman, H. M. (2001). The Forests of Taraba and Adamawa States, Nigeria. An Ecological Account and Plant Species Checklist (Vol. 57, p. 239). University of Canterbury. 10.2307/4110842

Chen, Y. Y., Satake, A., Sun, I. F., Kosugi, Y., Tani, M., Numata, S., Hubbell, S. P., Fletcher, C., Nur Supardi, M. N., & Wright, S. J. (2018). Species-specific flowering cues among general flowering Shorea species at the Pasoh Research Forest, Malaysia. Journal of Ecology, 106(2), 586–598. 10.1111/1365-2745.12836

Cleland, E. E., Chuine, I., Menzel, A., Mooney, H. A., & Schwartz, M. D. (2007). Shifting plant phenology in response to global change. Trends in Ecology and Evolution, 22(7), 357–365. 10.1016/j.tree.2007.04.003

Cleland, E. E., & Wolkovich, E. M. (2024). Effects of Phenology on Plant Community Assembly and Structure. Annual Review ofEcology, Evolution, and Systematics, 55, 471–492.

Condit, R., Watts, K., Bohlman, S. A., Pérez, R., Robin, B., Hubbell, S. P., Stephanie, A., Robin, B., & Stephen, P. (2002). Quantifying the Deciduousness of Tropical Forest Canopies under Varying Climates Stable URL : http://www.jstor.org/stable/3236572 REFERENCES Linked references are available on JSTOR for this article : Quantifying the deciduousness of tropical forest canop. Journal of Vegetation Science, 11(5), 649–658.

Corlett, R. T. (2016). The Impacts of Droughts in Tropical Forests. Trends in Plant Science, 21(7), 584–593. 10.1016/j.tplants.2016.02.003

Davis, C. C., Lyra, G. M., Park, D. S., Asprino, R., Maruyama, R., Torquato, D., Cook, B. I., & Ellison, A. M. (2022). New directions in tropical phenology. Trends in Ecology and Evolution, 37(8), 683–693. 10.1016/j.tree.2022.05.001

Edwards, W., Liddell, M. J., Franks, P., Nichols, C., & Laurance, S. G. W. (2018). Seasonal patterns in rainforest litterfall: Detecting endogenous and environmental influences from long-term sampling. Austral Ecology, 43(2), 225–235. 10.1111/aec.12559

Fidino, M., & Magle, S. B. (2017). Using Fourier series to estimate periodic patterns in dynamic occupancy models. Ecosphere, 8(9). 10.1002/ecs2.1944

Gouhier, T. C., & Guichard, F. (2014). Synchrony: Quantifying variability in space and time. Methods in Ecology and Evolution, 5(6), 524–533. 10.1111/2041-210X.12188

Hacket-Pain, A., Asenath Adienge, Bogdziewicz, M., Bush, E. R., Chapman, H., Memiaghe, H. R., Ofosu-Bamfo, B., Satake, A., Journé, V., & 13. (2024). Patterns of fruit production in tropical forests are shifting with negative outnumbering positive trends. EcoEvoRxiv. 10.32942/X2404C

Hacket-Pain, A., Foest, J. J., Pearse, I. S., LaMontagne, J. M., Koenig, W. D., Vacchiano, G., Bogdziewicz, M., Caignard, T., Celebias, P., Dormolen, J. van, Fernández-Martínez, M., Moris, J. V., Palaghianu, C., Pesendorfer, M., Satake, A., Schermer, E., Tanentzap, A. J., Thomas, P. A., Vecchio, D., … Ascoli, D. (2022). MASTREE+: Time-series of plant reproductive effort from six continents. Global Change Biology, 28(9), 3066–3082. 10.1111/gcb.16130

Ho, B. C., Chia, E. J. J., Chong, K. Y., Tan, J. S. Y., Tan, W. X., Lai, S., Choo, T. Y. S., Tan, P. Y., & Er, K. B. H. (2024). Changes in tropical leafing behaviour with climate change over nine decades: A case study from the Singapore Botanic Gardens. Plants People Planet. 10.1002/ppp3.10547

Iler, A. M., Caradonna, P. J., Forrest, J. R. K., & Post, E. (2021). Demographic Consequences of Phenological Shifts in Response to Climate Change. Annual Review of Ecology, Evolution, and Systematics, 52, 221–245. 10.1146/annurev-ecolsys-011921-032939

Janssen, T., Van Der Velde, Y., Hofhansl, F., Luyssaert, S., Naudts, K., Driessen, B., Fleischer, K., & Dolman, H. (2021). Drought effects on leaf fall, leaf flushing and stem growth in the Amazon forest: Reconciling remote sensing data and field observations. Biogeosciences, 18(14), 4445–4472. 10.5194/bg-18-4445-2021

Jeffs, C. T., Kennedy, P., Griffith, P., Gripenberg, S., Markesteijn, L., & Lewis, O. T. (2018). Seed predation by insects across a tropical forest precipitation gradient. Ecological Entomology, 43(6), 813–822. 10.1111/een.12672

Jenik, J., & Hall, J. B. (1966). The Ecological Effects of the Harmattan Wind in the Djebobo Massif (Togo Mountains, Ghana). The Journal of Ecology, 54(3), 767. 10.2307/2257816

Kuppler, J., & Kotowska, M. M. (2021). A meta-analysis of responses in floral traits and flower– visitor interactions to water deficit. Global Change Biology, 27(13), 3095–3108. 10.1111/gcb.15621

Laan-Luijkx, I. T. van der, Velde, I. R. van der, Krol, M. C., Gatti, L. V., Domingues, L. G., Correia, C. S. C., Miller, J. B., Gloor, M., Leeuwen, T. T. van, Kaiser, J. W., Wiedinmyer, C., Basu, S., Clerbaux, C., & Peters, W. (2015). Response of the Amazon carbon balance to the 2010 drought derived with CarbonTracker South America. Global Biogeochemical Cycles, 29, 1092–1108. 10.1111/1462-2920.13280

Lasky, J. R., Uriarte, M., & Muscarella, R. (2016). Synchrony, compensatory dynamics, and the functional trait basis of phenological diversity in a tropical dry forest tree community: Effects of rainfall seasonality. Environmental Research Letters, 11(11). 10.1088/1748-9326/11/11/115003

Lieberman, D., & Lieberman, M. (1984). The Causes and Consequences of Synchronous Flushing in a Dry Tropical Forest. Biotropica, 16(3), 193. 10.2307/2388052

Meir, P., & Pennington, R. T. (2011). Climatic change and seasonally dry tropical forests. In R. Dirzo, H. S. Young, H. A. Mooney, & G. Ceballos (Eds.), Seasonally dry tropical forests: Ecology and conservation (pp. 279–299). Island Press/Center for Resource Economics. 10.5822/978-1-61091-021-7_16

Mendoza, I., Condit, R. S., Wright, S. J., Caubère, A., Châtelet, P., Hardy, I., & Forget, P. M. (2018). Inter-annual variability of fruit timing and quantity at Nouragues (French Guiana): insights from hierarchical Bayesian analyses. Biotropica, 50(3), 431–441. 10.1111/btp.12560

Muñoz Sabater, J. (2019). ERA5-Land hourly data from 1950 to present. 10.24381/cds.e2161bac

Murphy, P. G., & Lugo, A. E. (1986). Ecology of tropical dry forest. Annual Review of Ecology and Systematics, 17(1986), 67–88. 10.1146/annurev.es.17.110186.000435

Nelson, W., Tong, Y. L., Lee, J. K., & Halberg, F. (1979). Methods for cosinor-rhythmometry. Chronobiologia, 6(4), 305–323.

Newstrom, L. E., Frankie, G. W., & Baker, H. G. (1994). A New Classification for Plant Phenology Based on Flowering Patterns in Lowland Tropical Rain Forest Trees at La Selva, Costa Rica. Biotropica, 26(2), 141. 10.2307/2388804

Nomura, N., Kikuzawa, K., & Kitayama, K. (2003). Leaf Flushing Phenology of Tropical Montane Rain Forests: Relationship to Soil Moisture and Nutrients. Tropics, 12(4), 261–276. 10.3759/tropics.12.261

Numata, S., Yamaguchi, K., Shimizu, M., Sakurai, G., Morimoto, A., Alias, N., Noor Azman, N. Z., Hosaka, T., & Satake, A. (2022). Impacts of climate change on reproductive phenology in tropical rainforests of Southeast Asia. Communications Biology, 5(1). 10.1038/s42003-022-03245-8

Ouédraogo, D. Y., Hardy, O. J., Doucet, J. L., Janssens, S. B., Wieringa, J. J., Stoffelen, P., Angoboy Ilondea, B., Baya, F., Beeckman, H., Daïnou, K., Dubiez, E., Gourlet-Fleury, S., & Fayolle, A. (2020). Latitudinal shift in the timing of flowering of tree species across tropical Africa: Insights from field observations and herbarium collections. Journal of Tropical Ecology. 10.1017/S0266467420000103

Ovaskainen, O., Skorokhodova, S., Yakovleva, M., Sukhov, A., Kutenkov, A., Kutenkova, N., Shcherbakov, A., Meyke, E., & Del Mar Delgado, M. (2013). Community-level phenological response to climate change. Proceedings of the National Academy of Sciences of the United States of America, 110(33), 13434–13439. 10.1073/pnas.1305533110

Pau, S., Wolkovich, E. M., Cook, B. I., Nytch, C. J., Regetz, J., Zimmerman, J. K., & Joseph Wright, S. (2013). Clouds and temperature drive dynamic changes in tropical flower production. Nature Climate Change, 3(9), 838–842. 10.1038/nclimate1934

R Core Team. (2022). R: A language and environment for statistical computing. R Foundation for Statistical Computing. https://www.R-project.org/

Reich, P. B., & Borchert, R. (1984). Water Stress and Tree Phenology in a Tropical Dry Forest in the Lowlands of Costa Rica. The Journal of Ecology, 72(1), 61. 10.2307/2260006

Renner, S. S., & Zohner, C. M. (2018). Climate change and phenological mismatch in trophic interactions among plants, insects, and vertebrates. Annual Review of Ecology, Evolution, and Systematics, 49, 165–182. 10.1146/annurev-ecolsys-110617-062535

Sakai, S. (2001). Phenological diversity in tropical forests. Population Ecology, 43(1), 77–86. 10.1007/PL00012018

Sakai, S., & Kitajima, K. (2019). Tropical phenology: Recent advances and perspectives. Ecological Research, 34(1), 50–54. 10.1111/1440-1703.1131

Sayer, E. J. (2006). Using experimental manipulation to assess the roles of leaf litter in the functioning of forest ecosystems. Biological Reviews, 81(1), 1–31. 10.1017/S1464793105006846

Sayer, E. J., Leitman, S. F., Wright, S. J., Rodtassana, C., Vincent, A. G., Bréchet, L. M., Castro, B., Lopez, O., Wallwork, A., & Tanner, E. V. J. (2024). Tropical forest above-ground productivity is maintained by nutrients cycled in litter. Journal of Ecology, December 2023, 1–11. 10.1111/1365-2745.14251

Shumway, R. H., & Stoffer, D. S. (2010). Time series analysis and its applications: with R examples. Springerlink.

Singh, K. P., & Kushwaha, C. P. (2016). Deciduousness in tropical trees and its potential as indicator of climate change: A review. Ecological Indicators, 69, 699–706. 10.1016/j.ecolind.2016.04.011

Siyum, Z. G. (2020). Tropical dry forest dynamics in the context of climate change: syntheses of drivers, gaps, and management perspectives. Ecological Processes, 9(1). 10.1186/s13717-020-00229-6

Stephenson, A. G. (1981). Flower and Fruit Abortion: Proximate Causes and Ultimate Functions. Annual Review of Ecology and Systematics, 12(1), 253–279. 10.1146/annurev.es.12.110181.001345

Sun, C., Kaplin, B. A., Kristensen, K. A., Munyaligoga, V., Mvukiyumwami, J., Kajondo, K. K., & Moermond, T. C. (1996). Tree Phenology in a Tropical Montane Forest in Rwanda. Biotropica, 28(4), 668–681.

Tang, J., Körner, C., Muraoka, H., Piao, S., Shen, M., Thackeray, S. J., & Yang, X. (2016). Emerging opportunities and challenges in phenology: A review. Ecosphere, 7(8), 1–17. 10.1002/ecs2.1436

Van Schaik, C. P., Terborgh, J. W., & Wright, S. J. (1993). The phenology of tropical forests: Adaptive significance and consequences for primary consumers. Annual Review of Ecology and Systematics, 24(1), 353–377. 10.1146/annurev.es.24.110193.002033

Wood, T. E., Lawrence, D., Clark, D. A., & Chazdon, R. L. (2009). Rain forest nutrient cycling and productivity in response to large-scale litter manipulation. Ecology, 90(1), 109–121. 10.1890/07-1146.1

Wright, S. J., & Calderón, O. (2006). Seasonal, El Niño and longer term changes in flower and seed production in a moist tropical forest. Ecology Letters, 9(1), 35–44. 10.1111/j.1461-0248.2005.00851.x

Wright, S. J., & Cornejo, F. H. (1990). Seasonal drought and leaf fall in a tropical forest. Ecology, 71(3), 1165–1175. 10.2307/1937384

Wu, J., Su, Y., Chen, X., Liu, L., Yang, X., Gong, F., Zhang, H., Xiong, X., & Zhang, D. (2021). Leaf shedding of Pan-Asian tropical evergreen forests depends on the synchrony of seasonal variations of rainfall and incoming solar radiation. Agricultural and Forest Meteorology, 311(October), 108691. 10.1016/j.agrformet.2021.108691

Wunderling, N., Staal, A., Sakschewski, B., Hirota, M., Tuinenburg, O. A., Donges, J. F., Barbosa, H. M. J., & Winkelmann, R. (2022). Recurrent droughts increase risk of cascading tipping events by outpacing adaptive capacities in the Amazon rainforest. Proceedings of the National Academy of Sciences of the United States of America, 119(32), 1–11. 10.1073/pnas.2120777119

Xu, C., McDowell, N. G., Fisher, R. A., Wei, L., Sevanto, S., Christoffersen, B. O., Weng, E., & Middleton, R. S. (2019). Increasing impacts of extreme droughts on vegetation productivity under climate change. Nature Climate Change, 9(12), 948–953. 10.1038/s41558-019-0630-6

Yadok, B. G., Forget, P. M., Gerhard, D., & Chapman, H. (2019). Low fruit-crop years of Carapa oreophila drive increased seed removal and predation by scatterhoarding rodents in a West African forest. Acta Oecologica, 99(February), 103448. 10.1016/j.actao.2019.103448

