## Supplementary information for "Changes in quantity and timing of foliar and reproductive phenology of tropical dry-forest trees under a warming and drying climate"

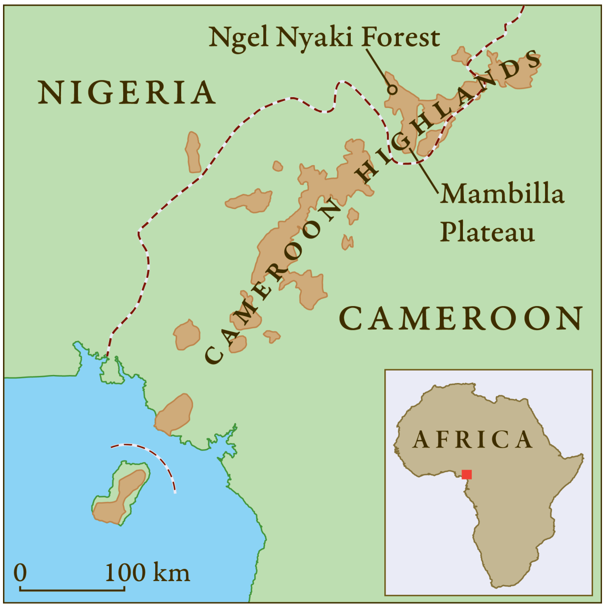


Figure S1: Map of the Cameroon Highlands showing the Mambilla Plateau and the location of Ngel Nyaki forest reserve. Modified from Thia (2014).

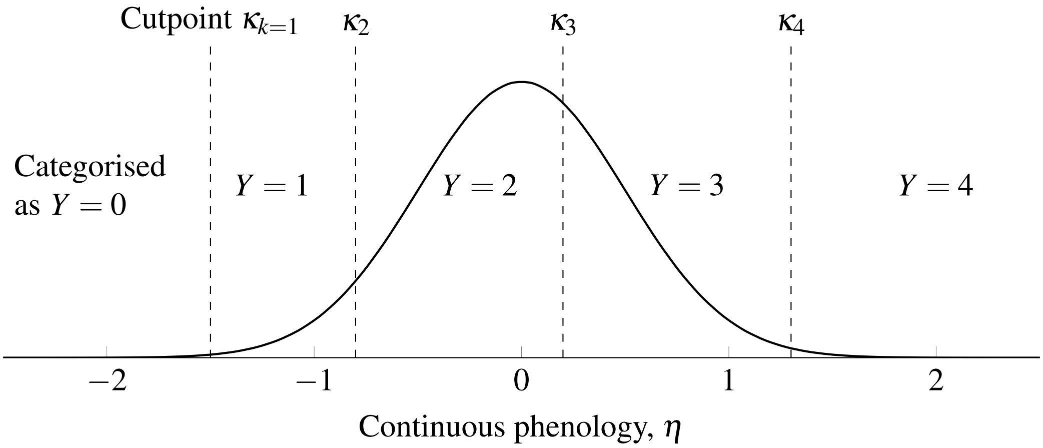


Figure S2: An illustrated explanation of the cumulative-logit regression (adapted from Bürkner and Vuorre 2019 for our phenology data), also known as ordered-logit regression (McElreath 2020). The model assumes that phenology is an underlying Normally-distributed, latent quantity $\eta$ that is hard to or cannot be measured by a field observer, who disretised the latent process into categorical, ordinal scores $Y$. To map the observed ordinal values to the underlying process, the model estimates (four) cutpoints $\kappa$ that partition the latent phenology into (five) observable categories.

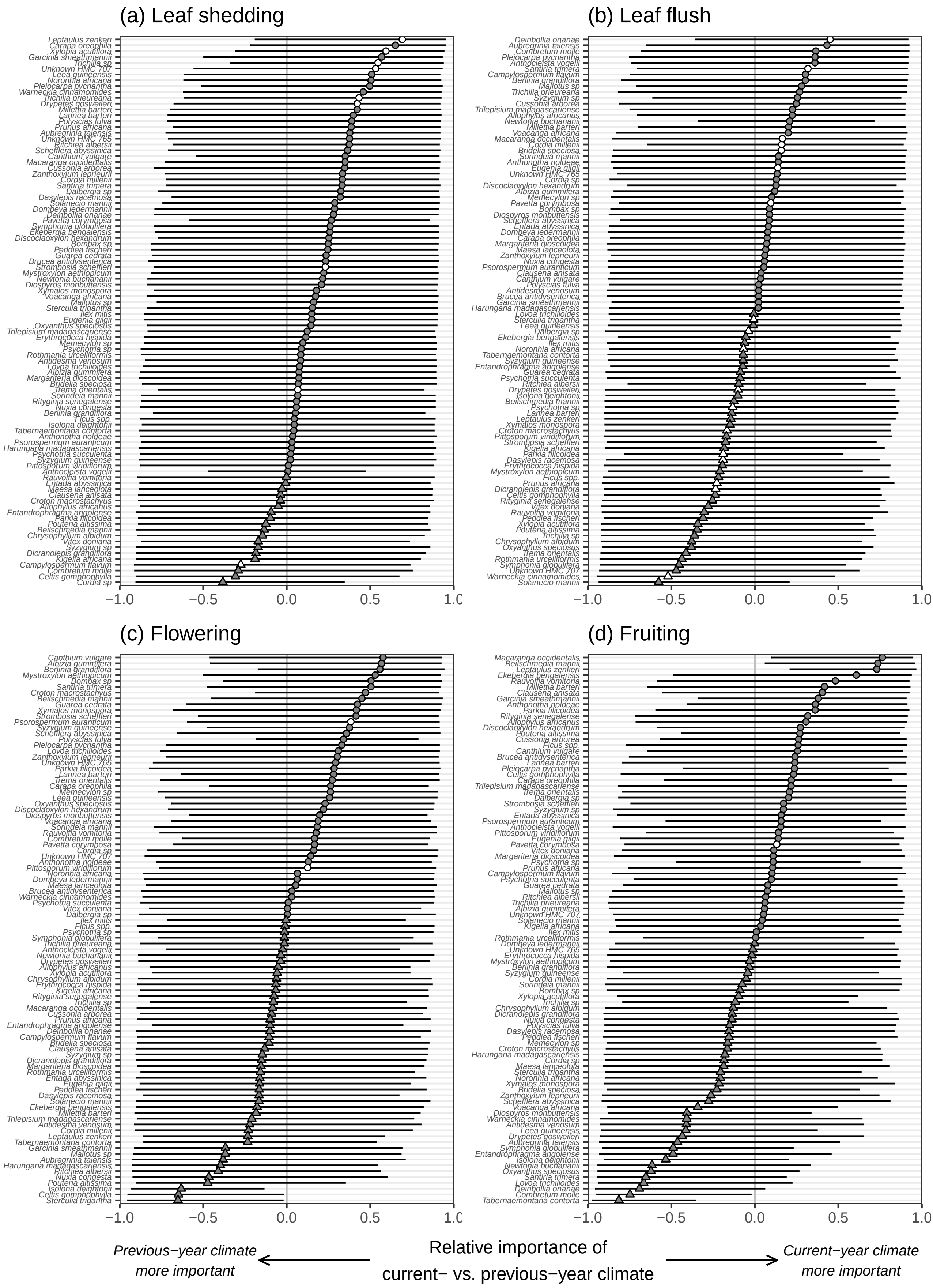


Figure S3: The relative importance of previous- vs. current-year climate variable on the phenology of species, calculated as the difference of the variance in phenology explained by each climate variable. The relative importance values range from $-1$ (all variance explained by previous-year climate) to 1 (all variance explained by current-year climate); zero indicates that previous- and current-year climate explained the same amount of variance in phenology. Points and error bars are posterior circular median and 89% credible intervals. Grey and white symbols represent species with predominantly annual and subannual phenology, respectively. Circles and triangles represent species that were more influenced by current- and previous-year climate variables, respectively.

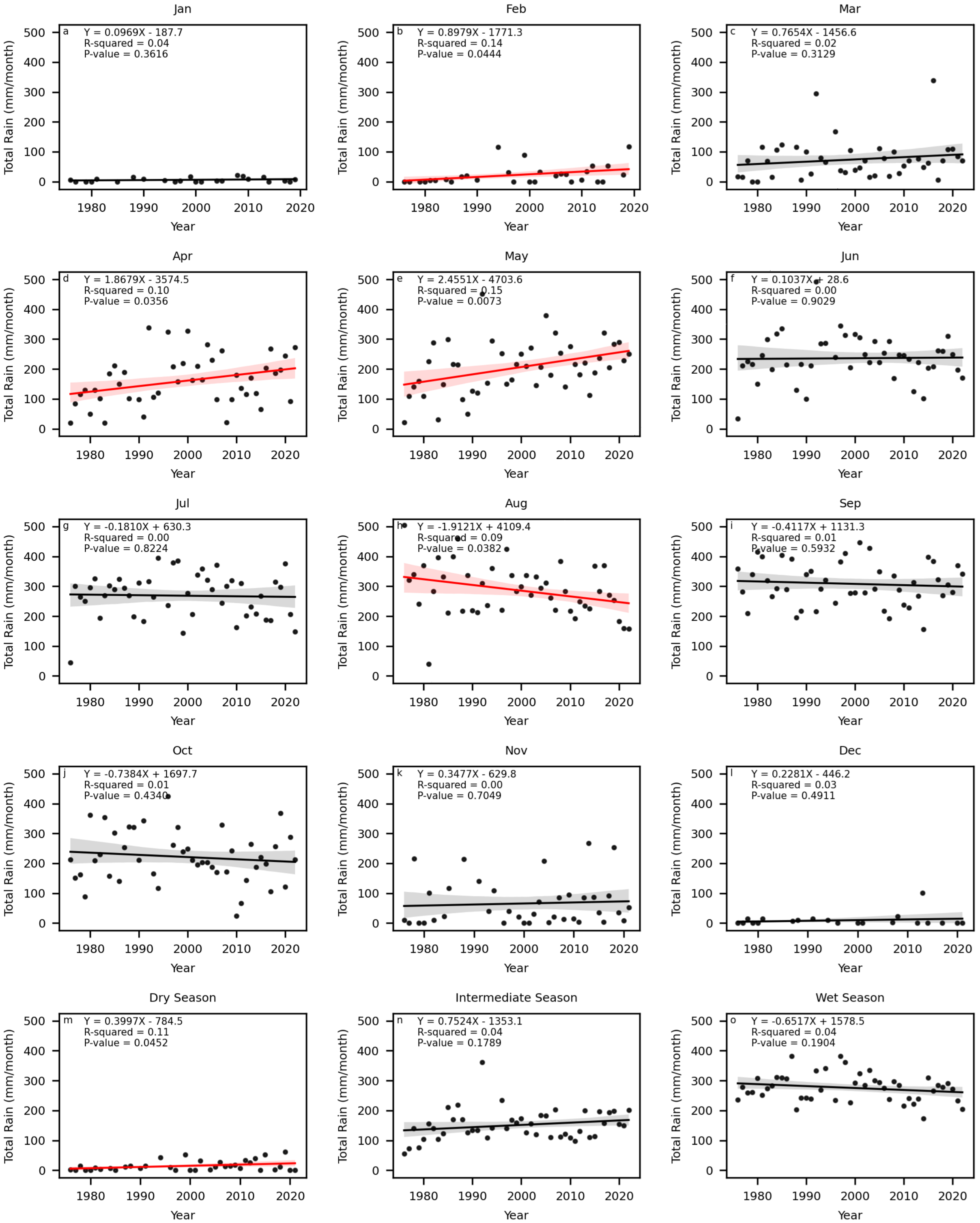


Figure S4: Monthly total rainfall measured at the Gembu weather station from 1976 to 2022. Panels **a** to **l** show individual months, and panels **m** to **o** seasonal groupings. The dry season includes December, January and February. The intermediate season includes March, April, May, October and November. The wet season includes June, July, August and September. Lines with shaded regions indicate linear regression with 90% confidence intervals (red and black colours denote $p$-value $\leq 0.1$ and otherwise, respectively). Fitted slopes are scaled to indicate per-year changes.

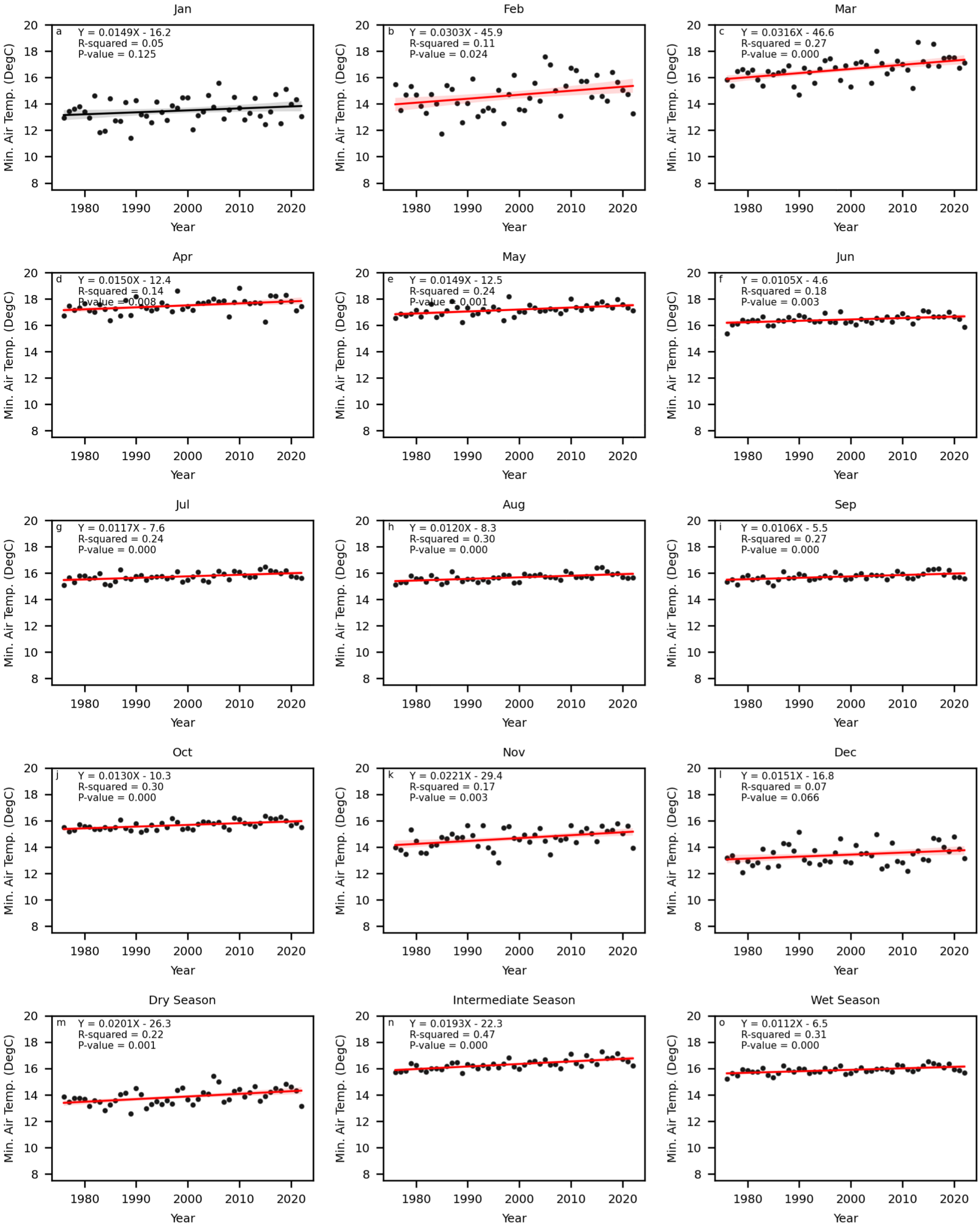


Figure S5: Monthly mean daily minimum air temperatures from the ERA5-reanalysis data product from 1976 to 2022. Panels **a** to **l** show individual months, and panels **m** to **o** seasonal groupings. The dry season includes December, January and February. The intermediate season includes March, April, May, October and November. The wet season includes June, July, August and September. Lines with shaded regions indicate linear regression with 90% confidence intervals (red and black colours denote $p$-value $\leq 0.1$ and otherwise, respectively). Fitted slopes are scaled to indicate per-year changes.

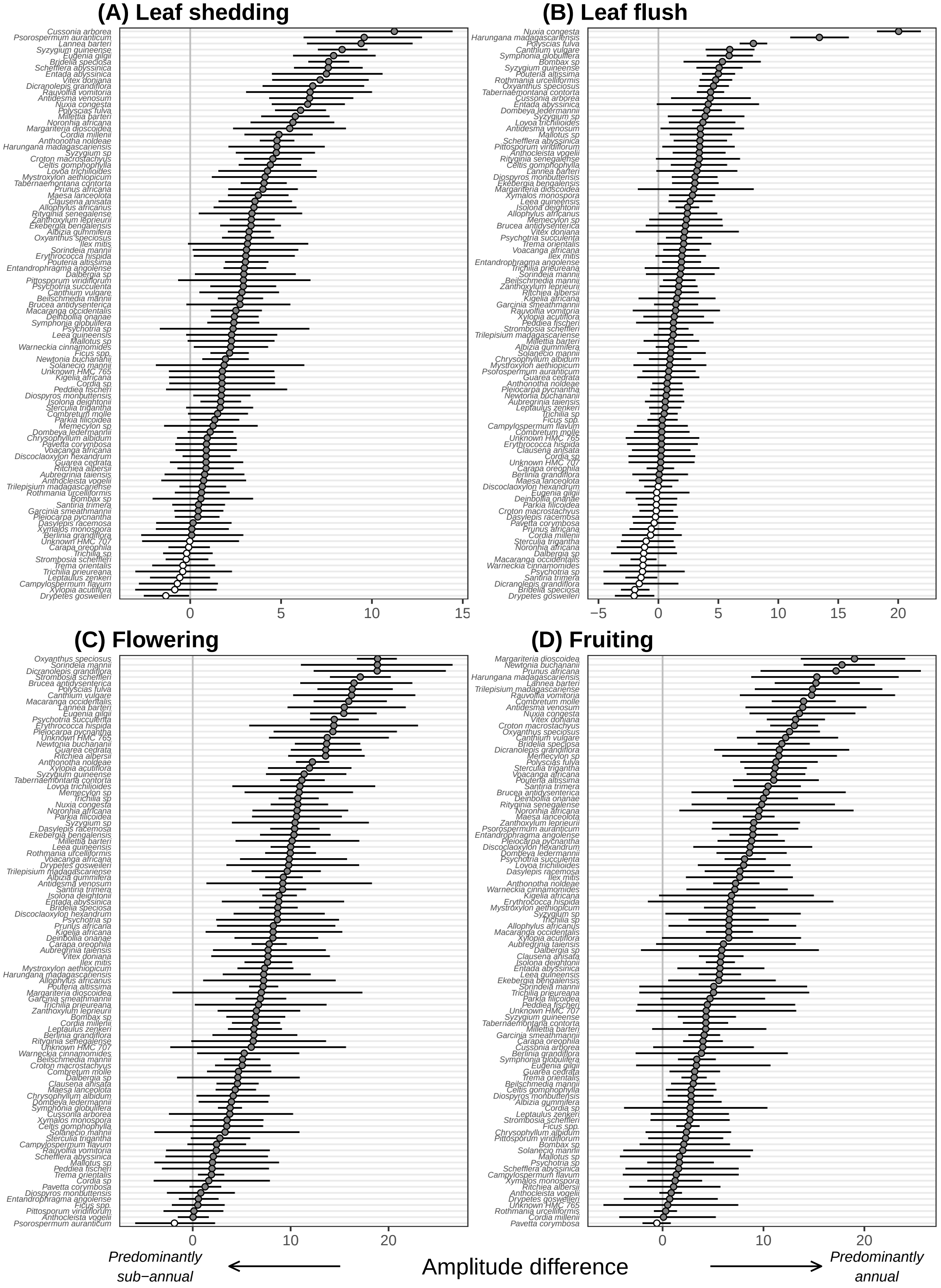


Figure S6: Species-specific difference between amplitudes of the two Fourier components. A positive value (grey symbols) indicates that the phenology of a species is predominantly annual (since we calculated amplitude of annual period − amplitude of sub-annual period), whereas a negative value (white symbols) indicates that a species is predominantly sub-annual. Points and error bars are posterior median and 89% credible intervals.

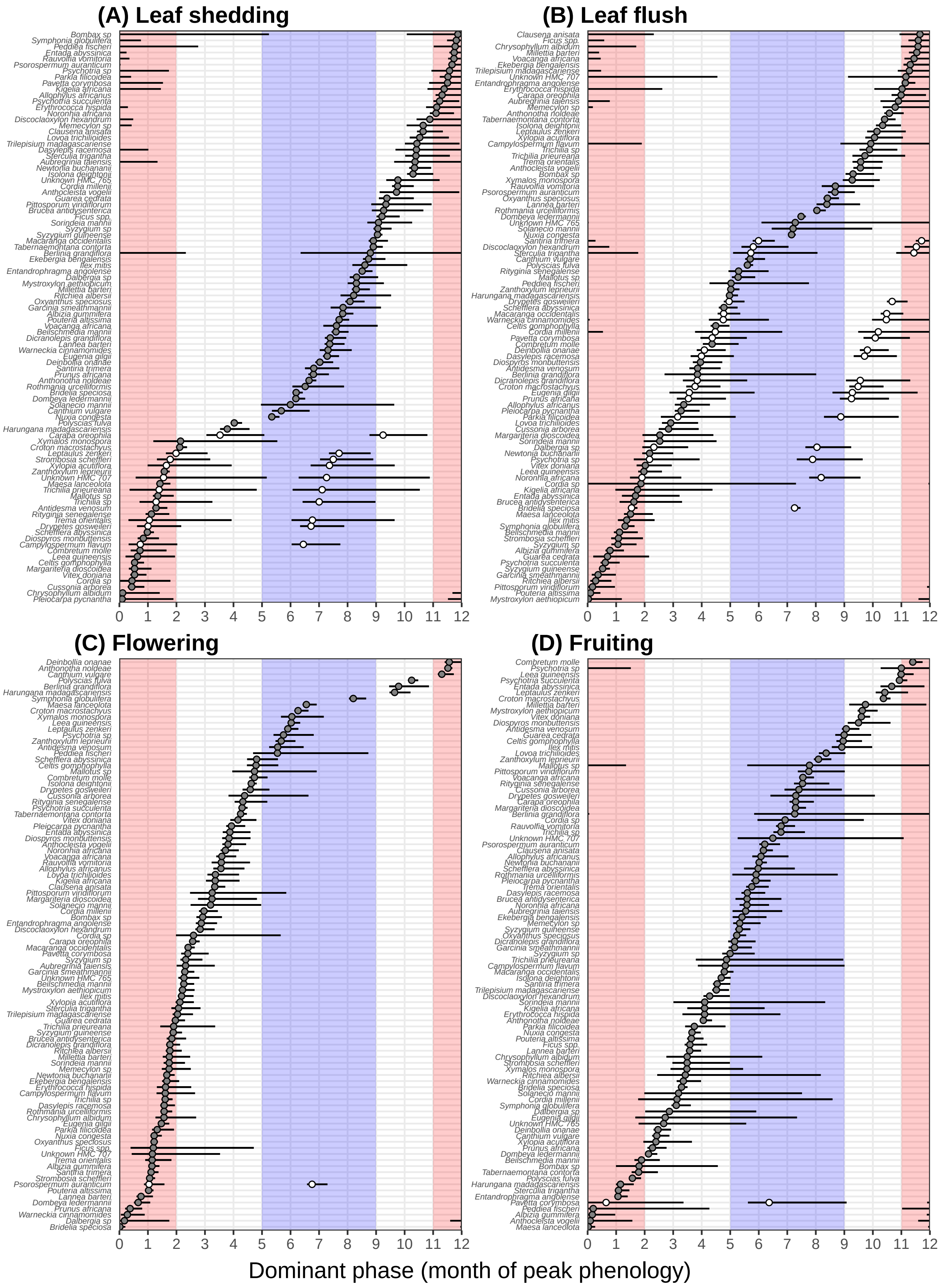


Figure S7: The phase (starting time of peak) of the dominant period of each species. The dominant period is taken as either annual (grey symbols) or sub-annual (white symbols) based on Fig. S6. Points and error bars are posterior circular median and 89% credible intervals. Blue and red shades denote wet and dry seasons. Note that sub-annual species have two phases because their phenologies occur twice per year.

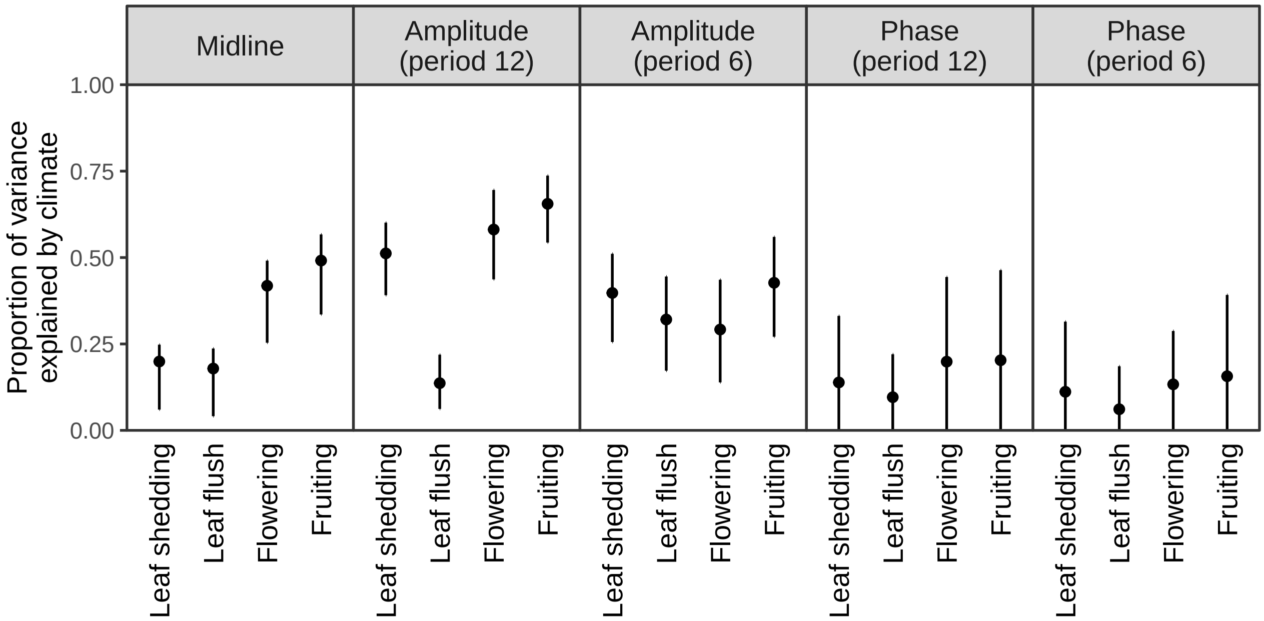


Figure S8: Proportion of variance in each periodic component of phenology explained by the climate variables. Points and error bars are posterior median and 89% credible intervals.

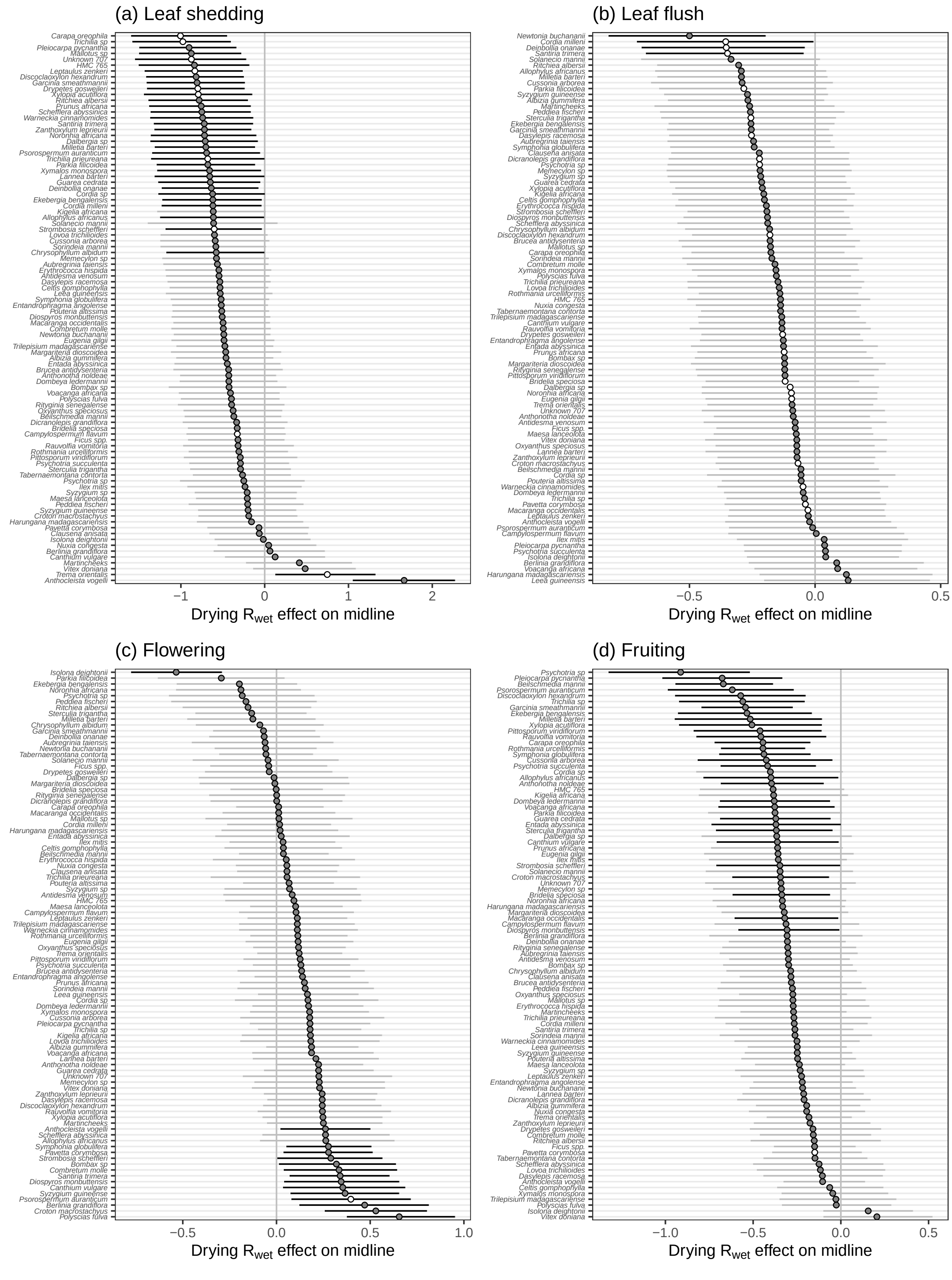


Figure S9: The effects of annual wet-season rainfall on the midline of leaf shedding **(a)**, leaf flush **(b)**, flowering **(c)**, and fruiting **(d)**. This figure is the same as the X-axes of Fig. 3's left column in the main text. Symbols and error bars are the posterior median and 89% credible intervals of species. Grey and white symbols represent species with predominantly annual and subannual phenology, respectively. Circles and triangles represent species that were more influenced by current- and previous-year climate variables, respectively.

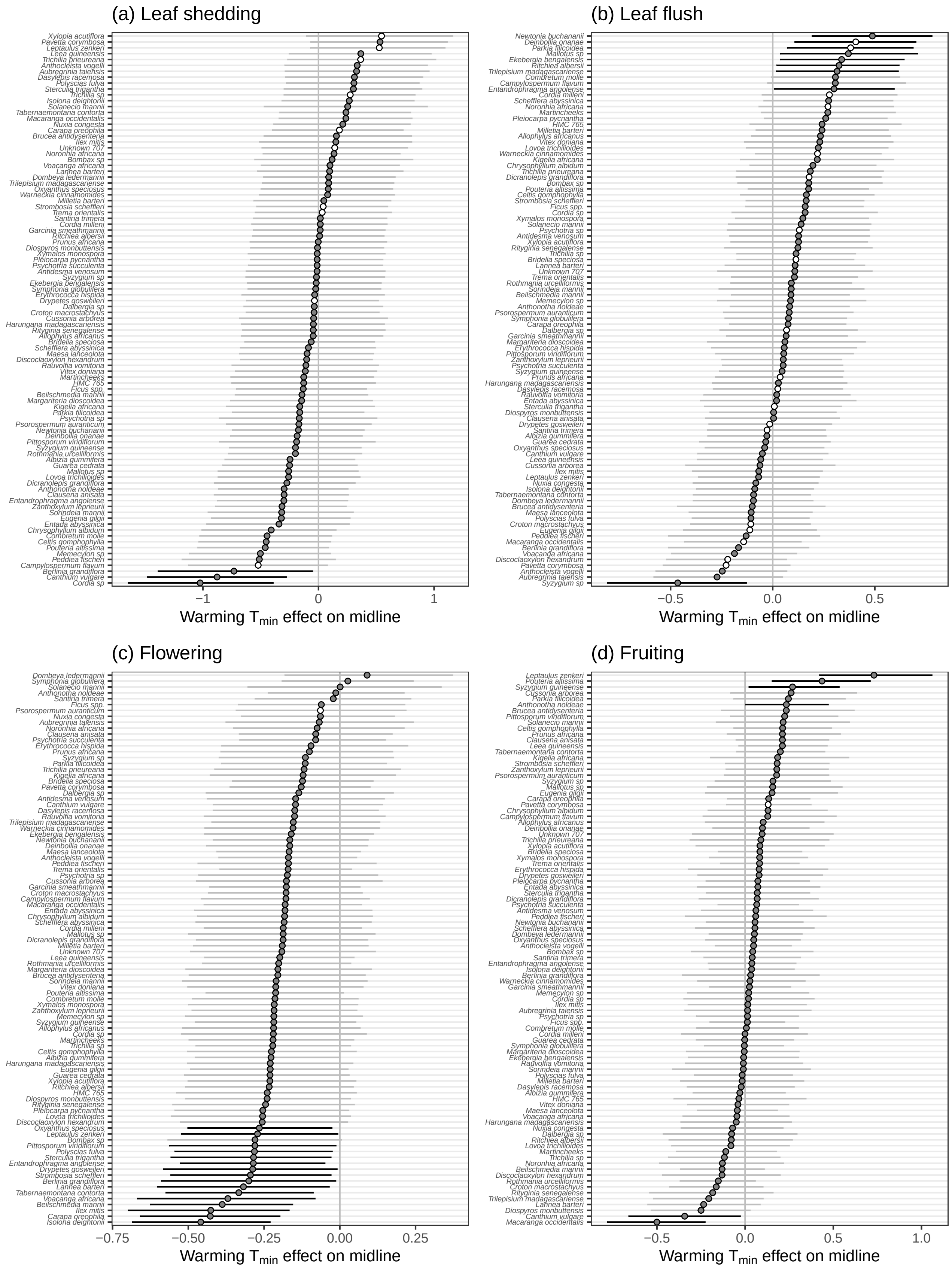


Figure S10: The effects of annual minimum temperature on the midline of leaf shedding **(a)**, leaf flush **(b)**, flowering **(c)**, and fruiting **(d)**. This figure is the same as the Y-axes of Fig. 3's left column in the main text. Symbols and error bars are the posterior median and 89% credible intervals of species. Grey and white symbols represent species with predominantly annual and subannual phenology, respectively. Circles and triangles represent species that were more influenced by current- and previous-year climate variables, respectively.

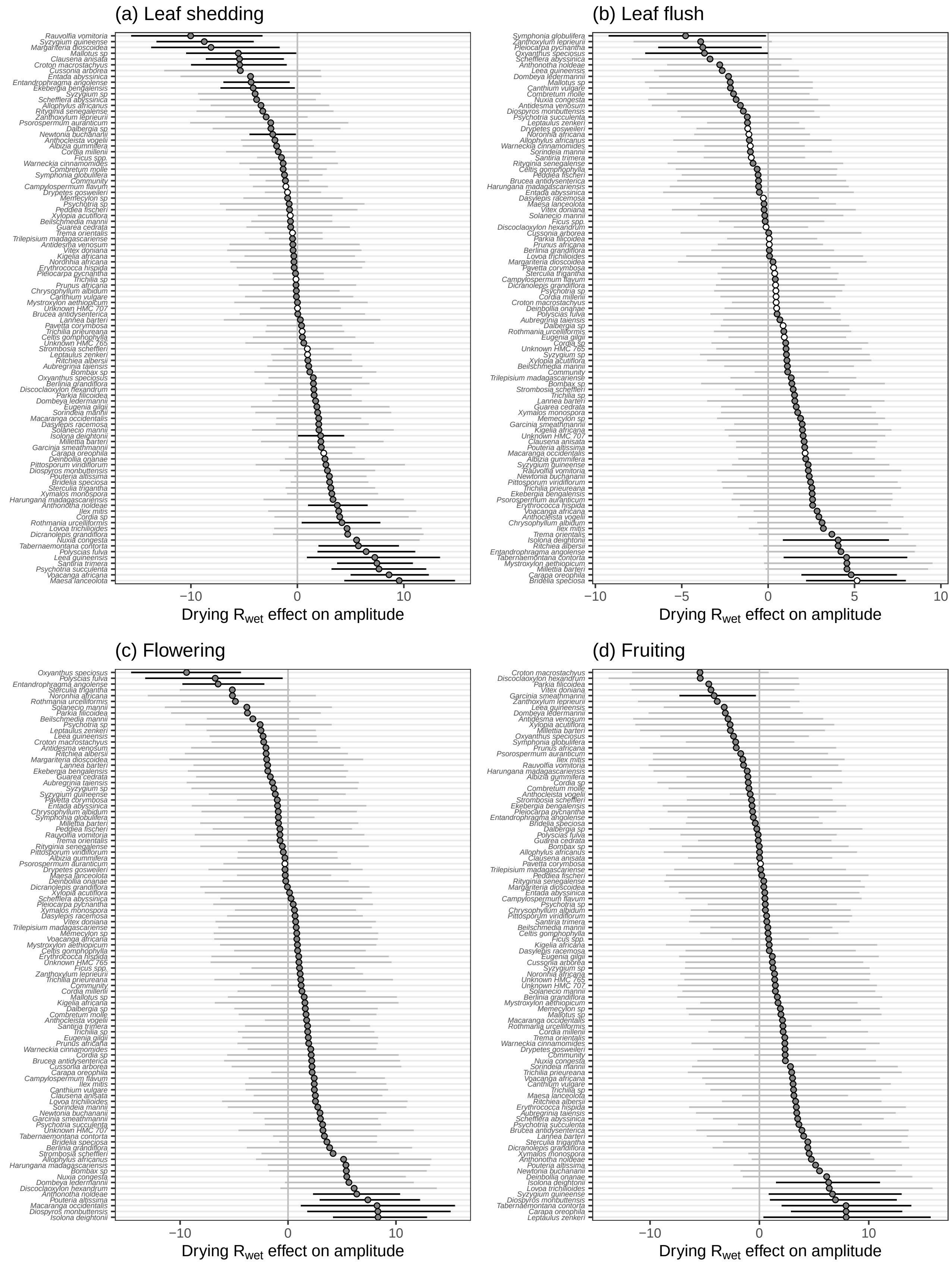


Figure S11: The effects of annual wet-season rainfall on the amplitude of leaf shedding **(a)**, leaf flush **(b)**, flowering **(c)**, and fruiting **(d)**. This figure is the same as the X-axes of Fig. 3's middle column in the main text. Symbols and error bars are the posterior median and 89% credible intervals of species. Grey and white symbols represent species with predominantly annual and subannual phenology, respectively. Circles and triangles represent species that were more influenced by current- and previous-year climate variables, respectively.

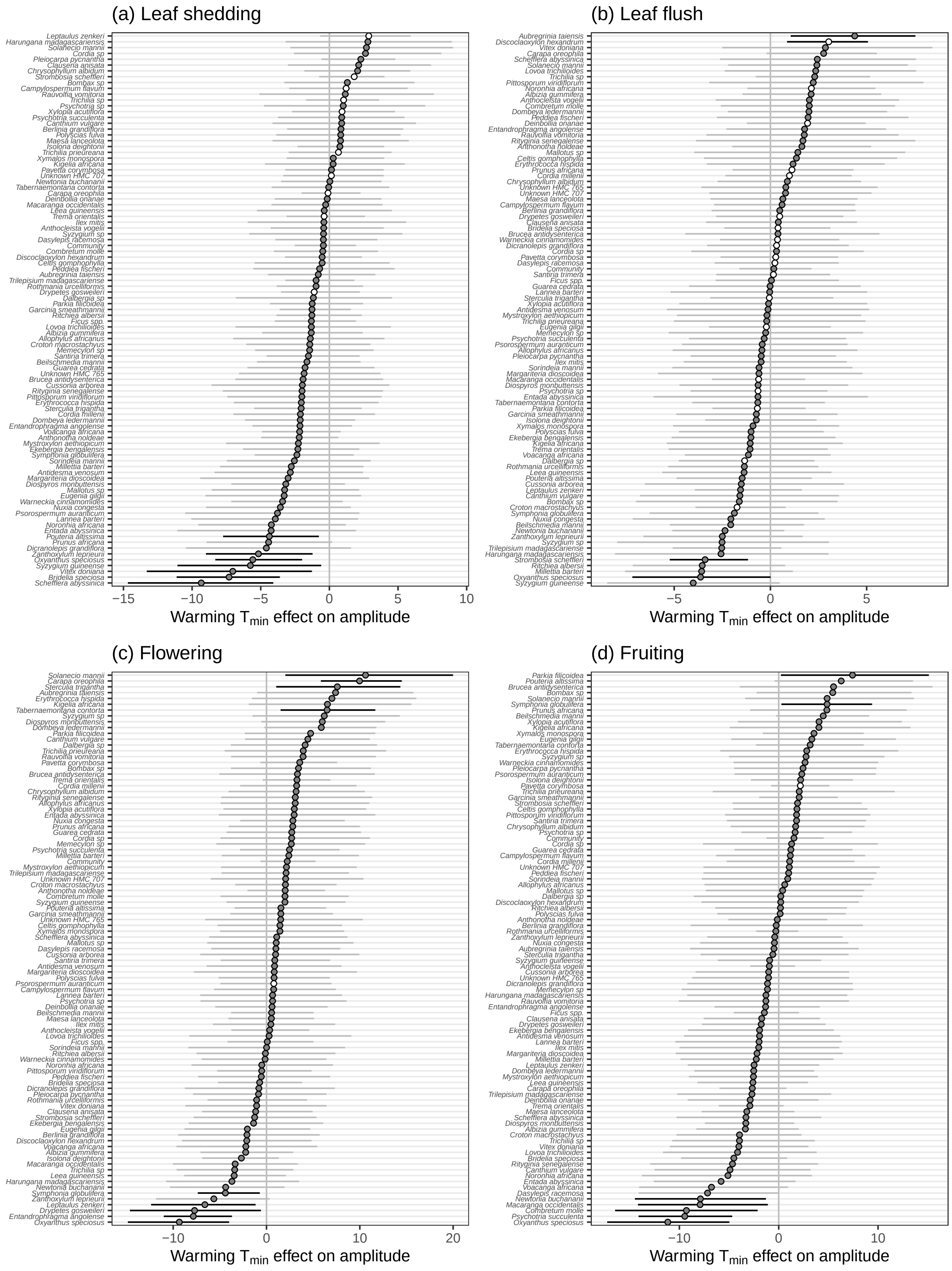


Figure S12: The effects of annual minimum temperature on the amplitude of leaf shedding **(a)**, leaf flush **(b)**, flowering **(c)**, and fruiting **(d)**. This figure is the same as the Y-axes of Fig. 3's middle column in the main text. Symbols and error bars are the posterior median and 89% credible intervals of species. Grey and white symbols represent species with predominantly annual and subannual phenology, respectively. Circles and triangles represent species that were more influenced by current- and previous-year climate variables, respectively.

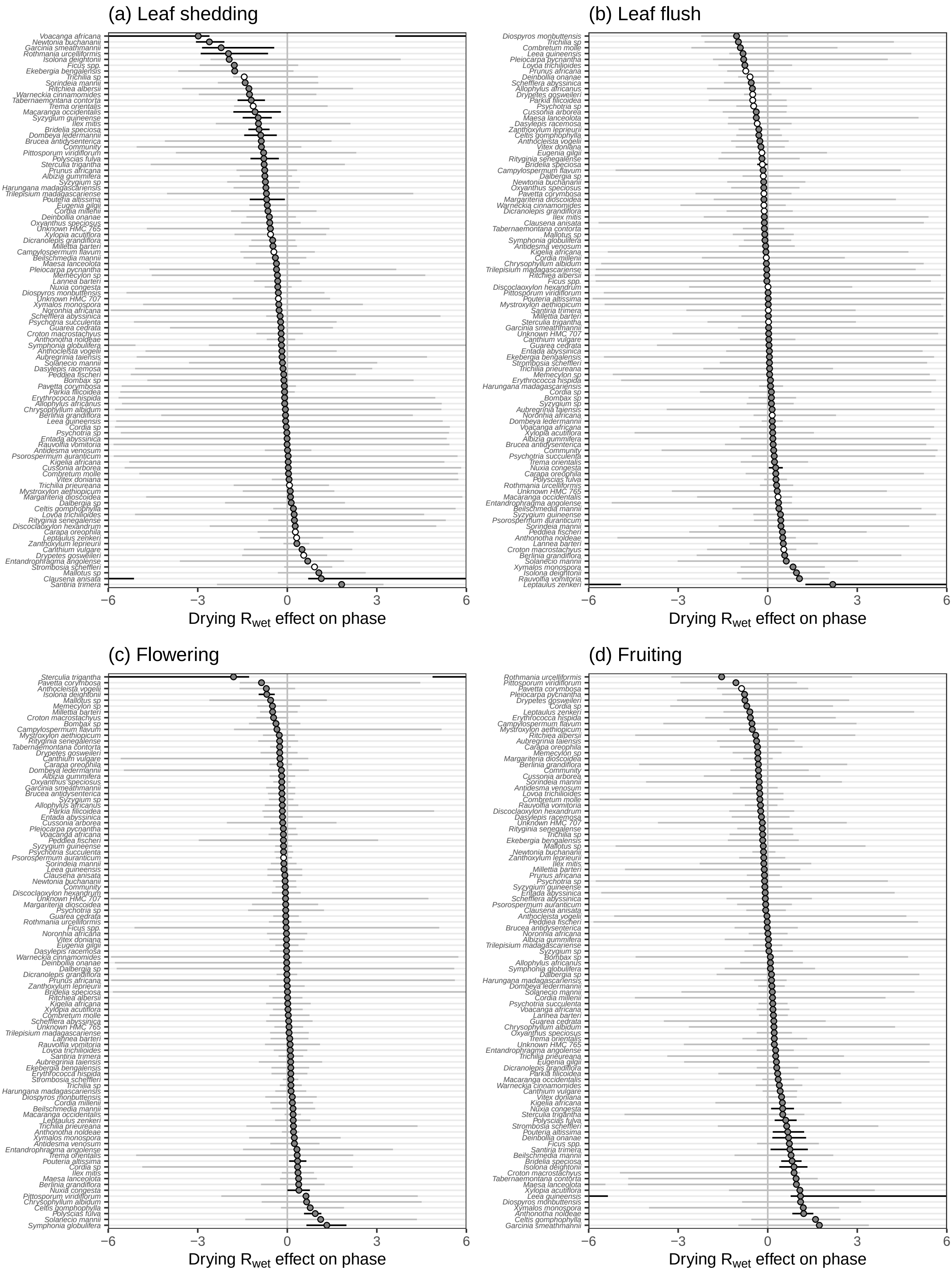


Figure S13: The effects of annual wet-season rainfall on the phase of leaf shedding **(a)**, leaf flush **(b)**, flowering **(c)**, and fruiting **(d)**. This figure is the same as the X-axes of Fig. 3's right column in the main text. Symbols and error bars are the posterior median and 89% credible intervals of species. Grey and white symbols represent species with predominantly annual and subannual phenology, respectively. Circles and triangles represent species that were more influenced by current- and previous-year climate variables, respectively.

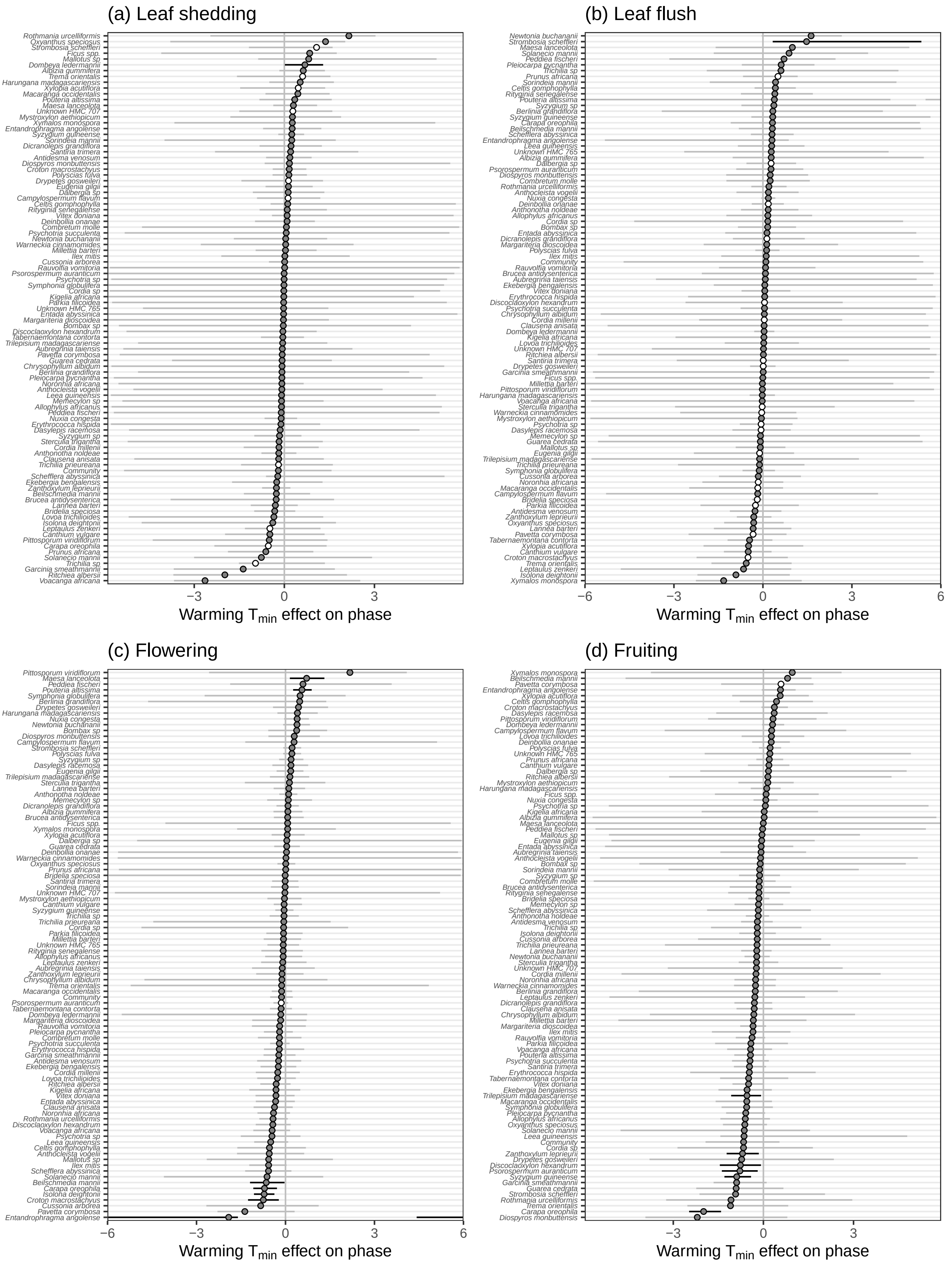


Figure S14: The effects of annual minimum temperature on the phase of leaf shedding **(a)**, leaf flush **(b)**, flowering **(c)**, and fruiting **(d)**. This figure is the same as the Y-axes of Fig. 3's right column in the main text. Symbols and error bars are the posterior median and 89% credible intervals of species. Grey and white symbols represent species with predominantly annual and subannual phenology, respectively. Circles and triangles represent species that were more influenced by current- and previous-year climate variables, respectively.

### S1 References

Bürkner, Paul Christian, and Matti Vuorre. 2019. “Ordinal Regression Models in Psychology: A Tutorial.” *Advances in Methods and Practices in Psychological Science* 2 (1): 77–101. https://doi.org/10.1177/2515245918823199.

McElreath, Richard. 2020. *Statistical rethinking: A bayesian course with examples in R and stan*. Second edi. CRC Press. https://doi.org/10.1201/9781315372495.

Thia, Joshua A. 2014. “The plight of trees in disturbed forest: conservation of Montane Trees, Nigeria.” MSc thesis, University of Canterbury.
